# Length-dependent Intramolecular Coil-to-Globule Transition in Poly(ADP-ribose) Induced by Cations

**DOI:** 10.1101/2023.10.25.564012

**Authors:** Tong Wang, Kush Coshic, Mohsen Badiee, Aleksei Aksimentiev, Lois Pollack, Anthony K. L. Leung

## Abstract

Poly(ADP-ribose) (PAR), a non-canonical nucleic acid, is essential for DNA/RNA metabolism and protein condensation, and its dysregulation is linked to cancer and neurodegeneration. However, key structural insights into PAR’s functions remain largely uncharacterized, hindered by the challenges in synthesizing and characterizing PAR, which are attributed to its length heterogeneity. A central issue is how PAR, comprised solely of ADP-ribose units, attains specificity in its binding and condensing proteins based on chain length. Here, we integrate molecular dynamics simulations with small-angle X-ray scattering to analyze PAR structures. We reveal the diverse structural ensembles of PAR and the factors influencing them, including a notable length-dependent compaction of PAR upon the addition of small amounts of Mg^2+^ ions. Unlike PAR_15_, PAR_22_ forms ADP-ribose bundles via local intramolecular coil-to-globule transitions. Understanding these length-dependent structural changes could be central to deciphering the specific biological functions of PAR.

## Introduction

Poly(ADP-ribose) (PAR) is a non-canonical nucleic acid produced in cells as a post-translational modification by ADP-ribosyltransferases commonly known as PARPs (Fig. 1A) ^1,2^. Functionally, PAR may serve as a signal mediator, conjugating to proteins in restoring homeostasis after stress, such as DNA damage and inflammation (Fig. 1B) ^3–7^. PAR may also serve as a scaffold for protein complex or biomolecular condensate formation, where its negative charge allows for multivalent noncovalent interactions with basic proteins ^8,9^.

**Figure 1.**
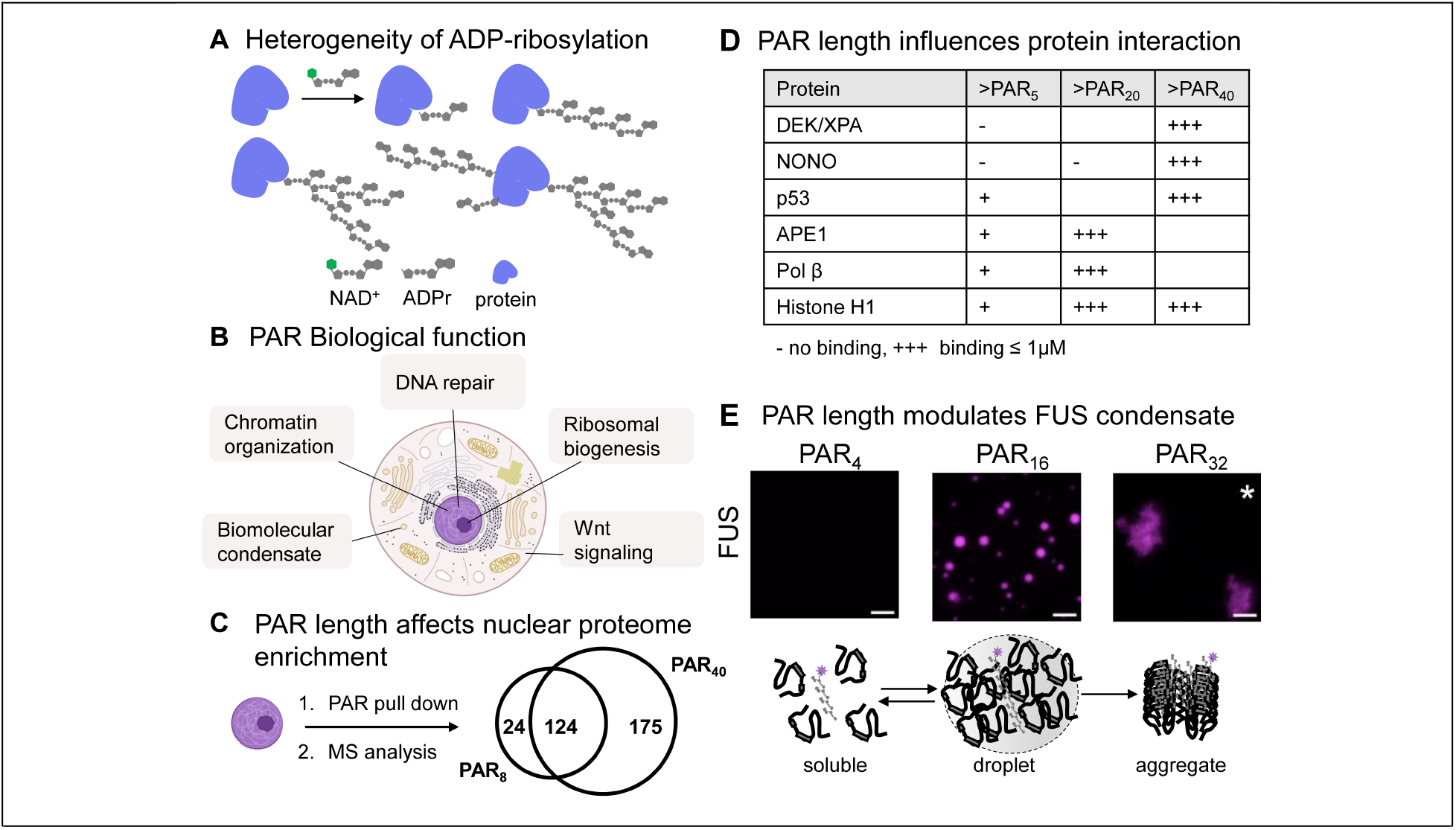
Biological functions of PAR and its relationship to chain length. **(A)** ADP-ribosylation is heterogenous, presented as monomeric, linear or branched polymer forms as poly(ADP-ribose) or PAR. **(B)** PAR mediates various cellular processes at distinct subcellular locations. **(C)** Pulldown of PAR-binding proteins in nuclear lysate coupled with mass spectrometry analysis indicates a preference of proteome to the length of PAR (data reproduced from Dasovich et al with permission)^12^. **(D)** Biochemical studies showing that the strength of PAR-proteins interaction is influenced by the chain length. **(E)** PAR modulates the formation and dynamics of biomolecular condensates linked to human disease. In vitro microscopic studies showing that PAR chain length influences material properties of FUS condensates, both at 1 µM; scale bar, 5 µm. These images were adopted from Rhine et al. with permission^16^.

The binding affinity of PAR to proteins depends on their chain length on a global scale, where its specificity could be even down to a single ADP-ribose difference (Fig. 1C, D) ^10–14^. In certain cases, such as with oncoprotein DEK, a threshold PAR length is necessary for appreciable binding to occur ^15^. Moreover, 4-mer PAR does not induce the condensation of FUS (a key protein associated with cancers and neurodegenerative diseases), whereas 8- and 16-mer do, and 32-mer drives its aggregation (Fig. 1E) ^16^. So, how does a simple homopolymer achieve such length-specific protein interactions? We posited that PAR may adopt distinct structures based on its polymer length.

Despite its functional importance, the 3D structure of PAR has not been extensively investigated, hampered by the challenges in synthesizing and characterizing PAR due to its heterogeneity in length. Previous circular dichroism analyses of mixed chain populations suggested the possibility of secondary structure formation contingent on cations ^17^. However, atomic structures beyond dimeric ADP-ribose have not been identified by experiments ^18–20^. Nuclear magnetic resonance studies did not identify any well-defined structures within populations characterized by mixed chain lengths ^21^. Molecular dynamics (MD) simulations reported multiglobular conformations in a 25-mer, but not in a 5-mer ^22^. This phenomenon occurs despite the rigidity of dihedral bonds at individual ribose-ribose linkages, where configurational entropy become predominant in longer polymers. Importantly, the simulations faced limitations in effectively sampling configurations attributed to their constrained duration (50 ns) and imperfections in the molecular force field describing cation-phosphate interactions ^23^. Notably, interpreting the dynamic structural ensembles of such macromolecules is complex, underscoring the need for advanced analysis and visualization methods. Although PAR’s functionality is influenced by its chain length, it remains uncertain whether a single structural model can capture the length-dependent diversity in biological PAR activity. A detailed analysis of the structural features of PAR at different lengths could elucidate its length-dependent conformations and potentially clarify its selective interactions with binding partners.

In mammalian cells under normal and mildly stressed conditions, PAR oligomers exhibit a size distribution spanning from as few as 2 units to approximately 20 units ^24–26^. Likewise, bacterial PARPs generate PAR within a similar range of lengths ^27,28^. Here, we integrated MD simulations with small-angle X-ray scattering (SAXS) measurements to determine structural ensembles of two physiologically relevant PAR lengths, PAR_15_ and PAR_22_, under various ionic conditions, yielding atomic-level snapshots of PAR structures substantiated by experimental data. This approach further distinguished itself from earlier MD analyses by incorporating multiple microsecond-long atomistic simulations, enhanced with the state-of-the-art corrections to nucleic acid force field parameters ^29^. This advance enabled us to accurately map the equilibrium ensembles of two PAR polymers. We further refined these conformational ensembles using SAXS data, thereby accurately assigning statistical weights to the conformations observed in the MD simulations. In addition, we employed graph theory as a systematic approach to categorize and visualize the PAR ensembles—a technique that could be broadly applied to the study of other disordered macromolecules.

Based on the SAXS-guided MD analyses, we defined structural order parameters to categorize backbone conformations across the structural ensembles of PAR_15_ and PAR_22_. Our results indicate that both backbone tortuosity and base stacking contribute to PAR compaction in unique ways for both lengths. By decomposing the structural ensembles into easily visualizable components using 3D class averaging in real space, we identified bundles of ADP-ribose units in PAR_22_, but not PAR_15_, in the presence of Mg^2+^. This detailed structural description offers a possible explanation for the heterogeneity in structural conformations and binding behaviors exhibited by PARs of different lengths.

## Results

### PAR Compaction and Structural Dynamics Influenced by Cationic Environment

To build a comprehensive understanding of PAR’s conformational behavior in different environments, we performed a series of MD simulations on PAR_22_. A typical simulation began with the PAR polymer in a fully extended state, submerged in a large volume of electrolyte solution (Fig. 2A). Within the first 50 ns of equilibration, the polymer transitioned to a more compact state (Fig. 2B). All conformations were characterized using the end-to-end distance (R_EE_) and the radius of gyration (R_g_) of the polymer.

**Figure 2.**
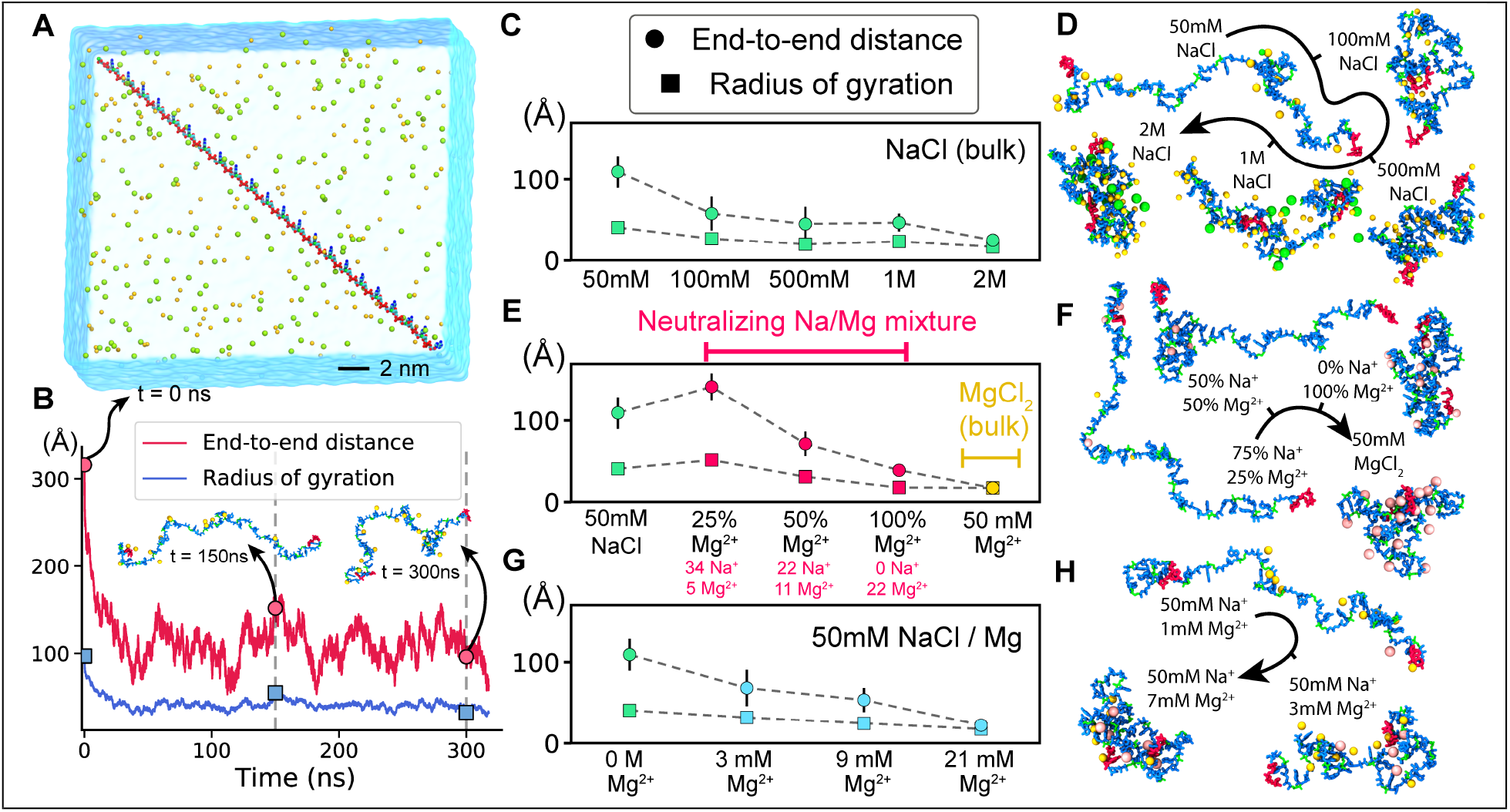
Molecular dynamics simulations of a 22-mer PAR polymer. **(A)** Initial configuration of a typical simulation system. One 22-mer PAR polymer is placed in electrolyte solution (semitransparent molecular surface) containing 50 mM of NaCl. **(B)** End-to-end distance (R_EE_) and radius of gyration (R_g_) of a PAR molecule as a function of time simulated in 50 mM NaCl electrolyte. **(C**,**E**,**G)** Average equilibrium end-to-end distance (circles) and radius of gyration (squares) of the 22-unit PAR polymer at various ion conditions. Dashed lines connect the points to guide the eye. Each data point represents a 250-ns trajectory average, after exclusion of the first 50 ns in each simulation where the molecule started in an extended state. **(D**,**F**,**H)** Representative snapshots of PAR conformation at the end of a 300 ns equilibration performed at the specified ion concentration conditions. The O3ʹ, C3ʹ, C4ʹ, C5ʹ, and O5ʹ atoms of PAR are shown in green whereas all other atoms in blue. Na^+^ (yellow), Cl^-^ (green) and Mg^2+^ (pink) ions located within 6 Å of PAR are shown as spheres. The ends of the PAR chains are depicted in red.

Prior research, including circular dichroism analyses of mixed-chain length PAR molecules and our recent single molecule Förster resonance energy transfer (smFRET) measurements, revealed that PAR compaction is sensitive to cations^17,30^. These findings align with established knowledge that nucleic acid structures are sensitive to their cationic environment^31,32^. In our MD simulations, we observed that both R_EE_ and R_g_ steadily decrease as the concentration of Na^+^ increases (Fig. 2C). Visualization of typical conformations indicates a transition: PAR adopts an extended, linear-like conformation at low salt concentrations and changes to a more compact, globular conformation at high salt concentrations (Fig. 2D). In the compact state, Na^+^ ions screened the backbone charge of PAR_22_, facilitating proximity between ADP-ribose units (Fig. 2D).

We extended our simulations to examine the impact of Mg^2+^ on PAR conformation and found an extreme sensitivity to this cation (Fig. 2E). In this set of simulations, we kept the total charge of cations equal in magnitude to the charge of PAR_22_, while varying the fraction of the charge neutralized by Mg^2+^ from 0 to 100% (Fig. 2E, red points). Remarkably, increasing the fraction of charge neutralization by Mg^2+^ from 25 to 50% led to a two-fold reduction in R_EE_. Further analysis revealed that Mg^2+^ ions do not compact PAR in a homogenous manner; instead, they induce the formation of highly compacted globules separated by extended polymer chains (Fig. 2F). This compaction was visualized over time across different systems using heatmaps that depict the number of neighboring ADP-ribose units within a 10 Å radius of each PAR residue (Fig. S1). Counting the net charge within this radius over time further revealed the interplay between spatial localization and charge (Fig. S2). Interestingly, the globular conformations observed at a neutralizing concentration of Mg^2+^ (100%, 7 mM) closely resemble those at much higher MgCl_2_ concentrations (50 mM). These data indicate that PAR_22_ is optimized for compaction even at Mg^2+^ concentrations much closer to physiological levels. This high sensitivity of PAR to Mg^2+^, including the formation of locally compacted domains (Movie S1 and S2), suggests that Mg^2+^ may play a structural role. Beyond simply screening electrostatic charges, Mg^2+^ may trigger a transition of PAR from an elongated polymer to a condensed globule.

To further quantify the effect of adding Mg^2+^, we conducted simulations with a fixed NaCl concentration of 50 mM while varying the MgCl_2_ concentration (Fig. 2G). These conditions align with those previously examined through smFRET experiments^30^. Our simulation showed that, in the presence of Mg^2+^, PAR transitioned from extended to compacted states and exhibited conformations where both states coexist within the same molecule (Fig. 2H, S1, S2). When compared to simulations in pure NaCl solvent (Fig. 2C-D), the onset of compact conformations occurred at significantly lower MgCl_2_ concentrations. Specifically, a complete globular collapse occurred at 21 mM of MgCl_2_, in contrast to 2 M NaCl. Furthermore, over 39% of total compaction was achieved at a MgCl_2_ concentration as low as 3 mM. These observations are consistent with previous studies on single-stranded RNA, where only 5 mM MgCl_2_ was required to induce the same R_g_ change as 600 mM NaCl ^31^.

Taken together, our MD simulations show that PAR is structurally dynamic, adopting a range of conformations depending on the ionic environment. In the absence of Mg^2+^, PAR adopts extended conformations at physiological Na^+^ concentrations. However, even small amounts of Mg^2+^ can trigger local compaction of the PAR polymer—a structural transition that we further explored in the remainder of this study.

### SAXS Reveals Distinct Compaction for PAR_15_ and PAR_22_ with Mg^2+^

Having surveyed a broad range of ionic conditions in the MD simulations of PAR, we focused on specific conditions for experimentally identifying various structural parameters of PAR using SAXS. We examined PAR_15_ and PAR_22_, both of which are found in normal and cancer cells, with the shorter one being more abundant ^24–26^. The SAXS experiments were conducted in a 100 mM NaCl environment to approach physiological conditions, and we assessed the impact of adding 1 mM MgCl_2_ on PAR compaction.

SAXS provides us with the overall size of the PAR structural ensembles, represented by the R_g_ values (Fig. 3A-B) ^33^. In a 100 mM NaCl solution, PAR_15_ had an R_g_ of 24.4 ± 1.2 Å, while PAR_22_ had an R_g_ of 32.6 ± 0.5 Å (Fig. 3B). Adding 1 mM Mg^2+^ led to a 5.3% reduction in R_g_ for PAR_15_ and a significant 19.0% reduction for PAR22, indicating a greater compacting effect on the longer PAR_22_ molecule.

**Figure 3.**
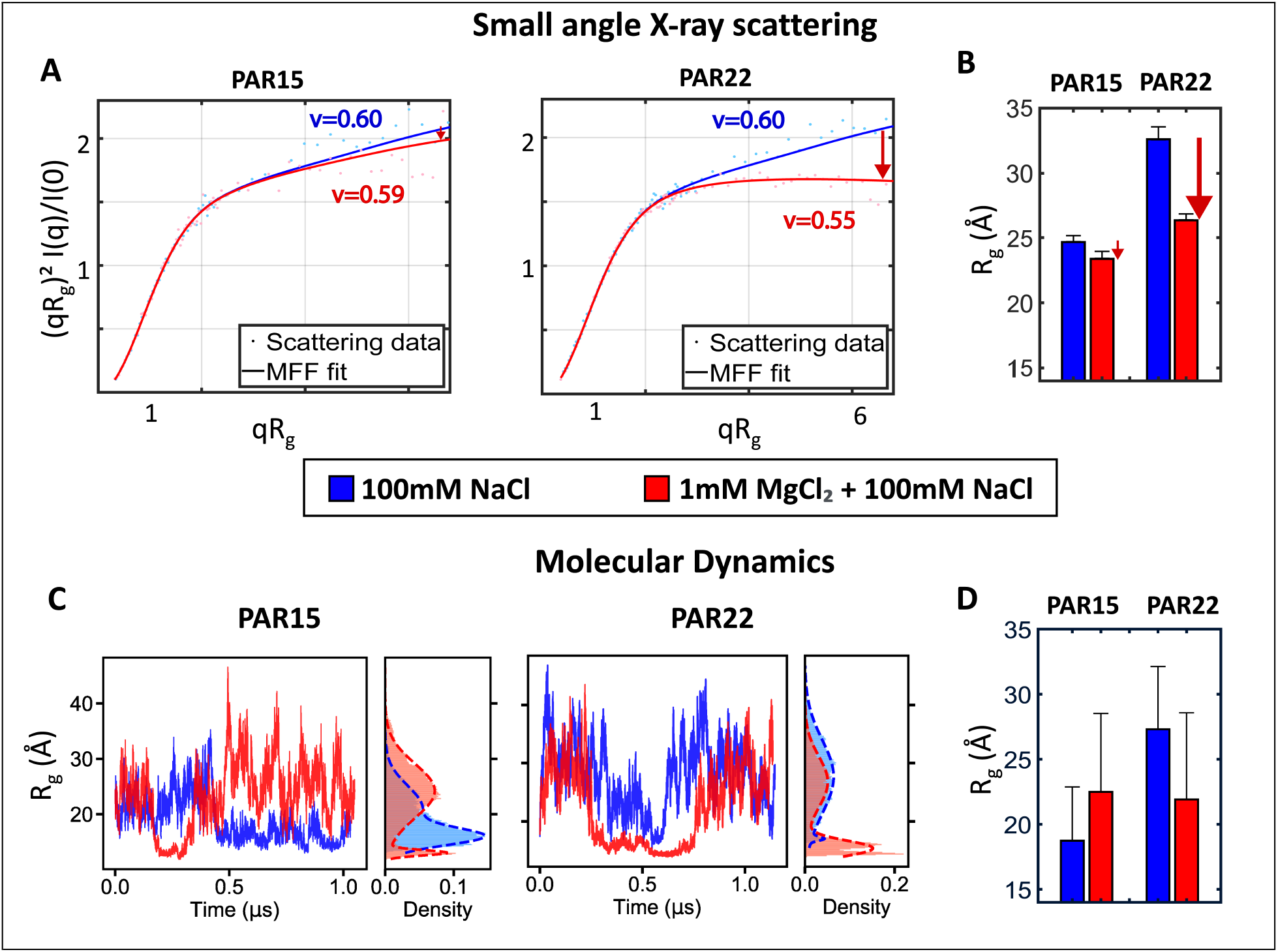
Length-dependent collapse of PAR polymer. **(A)** SAXS profiles of PAR_15_ and PAR_22_ in 100 mM NaCl with (red) and without (blue) the addition of 1 mM MgCl_2_. Data are plotted in dimensionless Kratky axes, normalizing out size differences and emphasizing changes in shape and disorder in the mid-angle scattering regime. Experimental scattering is shown in light colored points and solid lines show molecular form factor (MFF) model fits to the data, extracting the Flory scaling parameter ν (Fig. S3) ^34^. Reduced χ^2^ values for the fits are: 0.129 & 0.168 for PAR_15_ and 0.421 & 0.369 for PAR_22,_ without and with MgCl_2_ respectively. **(B)** SAXS-derived R_g_ values for PAR_15_ and PAR_22_ in the conditions assayed. Error bars show errors in the linear Guinier fits used to extract R_g_. (Fig. S4) **(C)** Radius of gyration of PAR_15_ and PAR_22_ polymers in MD simulations carried out at 100 mM NaCl, with and without 1 mM MgCl_2_. The histograms next to the timeseries plots illustrate the distribution of the R_g_ values. **(D)** Average simulated radius of gyration of PAR_15_ and PAR_22_ determined as a weighted mean of the two-Gaussian fit to the histograms.

We also calculated the Flory (ν) parameters for both PAR lengths at each salt concentration to understand their interaction with its surrounding solvent (Fig. 3A, S3) ^34^. Kratky plots were used for clearer data visualization, emphasizing the mid-angle scattering regime where the overall shape and degree of disorder of the molecular ensemble can be discerned ^35^.

When Mg^2+^ was absent, fits to SAXS profiles for both PAR_15_ and PAR_22_ yielded a ν value of 0.60 ± 0.02 and 0.60 ± 0.01, respectively. This ν value is expected for a self-avoiding random walk, suggesting that similar polymer properties for both PAR lengths in a 100 mM Na^+^ environment. The ratio of R_g_ values between PAR_22_ and PAR_15_ (32.6 Å / 24.4 Å = 1.34) is consistent with the expected behavior for molecules with ν = 0.6, according to the classical scaling law, R_g_ α length^v^. This agreement provides additional confidence in this polymer description of PAR.

When Mg^2+^ was introduced, both PAR lengths underwent a decrease in ν, more pronounced for PAR_22_ (ν = 0.55 ± 0.01) than for PAR_15_ (ν = 0.59 ± 0.02). These changes point to a stronger progression of PAR_22_ toward a theta state, where no interactions are present between polymer and solvent (Fig. 3A, S3) ^36^. Overall, these experimental results indicate that the longer PAR_22_ undergoes more significant conformational change when exposed to Mg^2+^ compared to its shorter counterpart, PAR_15_ (Fig. 3B).

### MD-SAXS Reveals Mg^2+^ Increases Tortuosity and Base Stacking in PAR_22_ Than PAR_15_

To examine specific conformations in PAR responsible for the observed differences in size and shape, we integrated SAXS data with additional MD simulations. We extended the simulations to 1 µs to capture a broader range of conformations for both PAR lengths. During these simulations, we noticed that the PAR conformations oscillated between extended and compact conformations (Fig. 3C). Because of this bimodal behavior, and since the duration of simulation duration was comparable to the lifetime of each state, we were unable to determine average R_g_ values for direct comparison (*i*.*e*., a large error margin when averaging R_g_ values across the entire simulations; Fig. 3D). Instead, we employed an ensemble optimization method (EOM) to refine the full pool of MD structures using SAXS data ^37,38^. The computed scattering profiles from the refined ensembles closely matched the experimentally measured SAXS profiles in molecular shapes and overall R_g_ values (Fig. 4A-B, S5,6).

**Figure 4.**
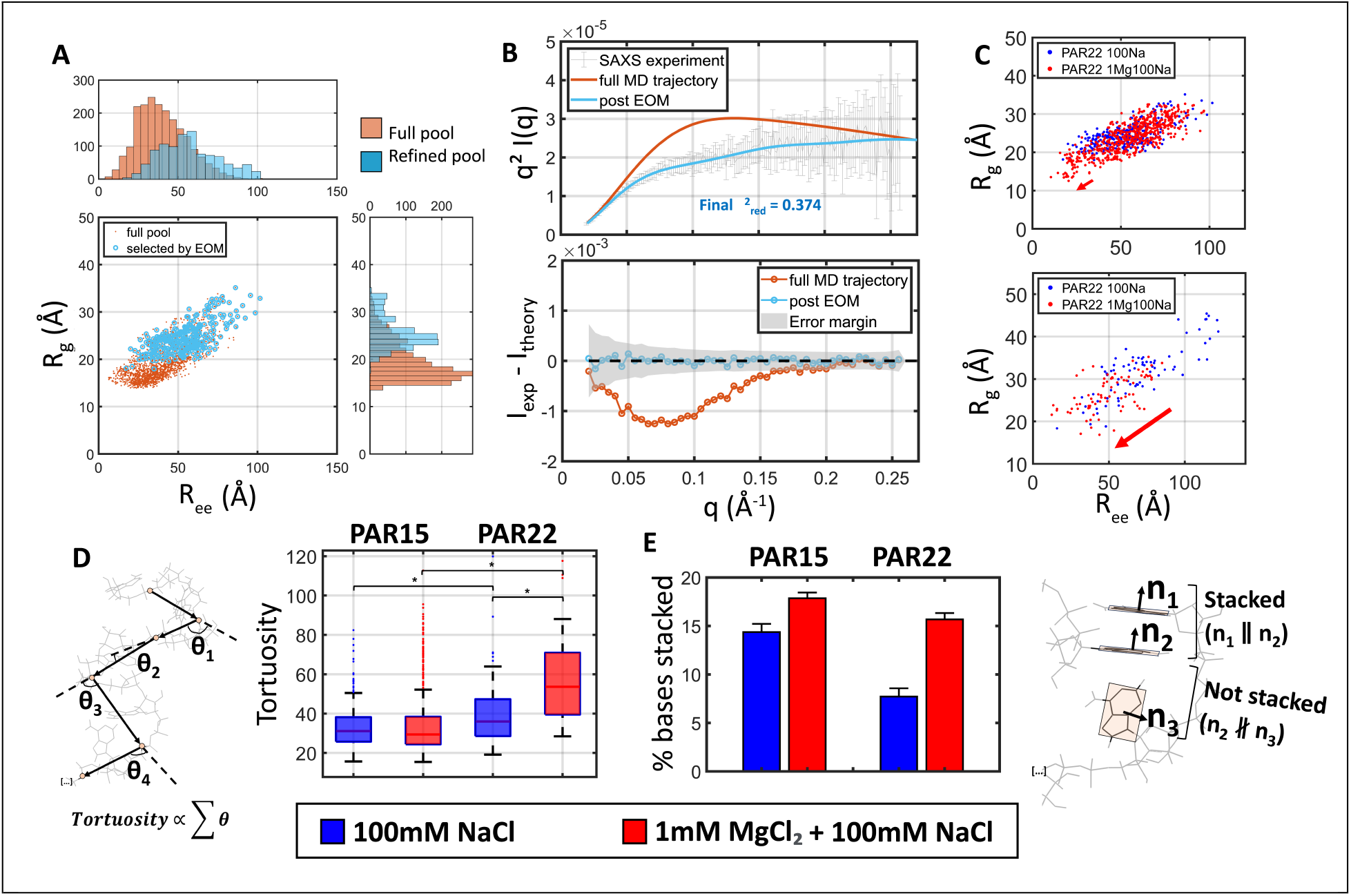
Determining structural ensembles for PAR using MD and SAXS. As an example, the case for PAR_15_ in 100 mM NaCl is shown. The same plots for PAR_22_ are shown in Fig. S5-6. **(A)** The pool of structures from the entire MD simulation is shown in orange, and the subset ensemble that agrees with the SAXS data in blue. Structures are parameterized in {R_g_,R_EE_} space. 1D histograms are weighted by the prevalence of each structure in the final ensembles. **(B)** Final agreement of the structural ensemble determined by EOM to the SAXS data, compared to initial agreement of the structural ensemble of all MD conformers. Residuals are shown in the bottom plot. **(C)** Ensembles of EOM-determined PAR structures, with and without Mg^2+^ for PAR_15_ and PAR_22_. Arrows show differential shifts to more compact states with the addition of Mg^2+^. **(D)** Tortuosities and **(E)** Fraction of adenine bases that are stacked in each structural ensemble of PAR. For the tortuosity plot, ‘*’ denotes p<0.05. For the base stacking plot, error bars show standard error.

Next, we analyzed how PAR conformation and compaction change with Mg^2+^ for both PAR_15_ and PAR_22_ (Fig. 4C). The refined pools displayed a relatively uniform distribution of structures around an average, with no pronounced bimodality, as one would expect for a macromolecular ensemble (Fig. S5). Importantly, the final fits between the EOM structural ensembles and SAXS data consistently fell within the experimental error margin, with ensemble R_g_ values in agreement across both methods (Fig. S6).

With refined ensembles now available for all conditions (PAR_15_ and PAR_22_, both with and without Mg^2+^), we analyzed the included structures. We computed the tortuosity index of each structure in the identified ensemble to gauge their backbone conformations (Fig. 4D). Tortuosity quantifies how “twisted” the polymer is compared to a straight line drawn connecting its endpoints. Without Mg^2+^, PAR_22_ has a significantly greater mean tortuosity across its structural ensemble than PAR_15_, indicating a more twisted backbone (Fig. 4D), despite the identical v values. Interestingly, introducing Mg^2+^ significantly increased the tortuosity of PAR_22_, but not PAR_15_ (Fig. 4D).

We also examined the role of π-π stacking in driving PAR chain compaction (Fig. 4E). Such interactions are particularly common among adenine bases and are known to induce intra-chain helicity in adenine-rich RNA sequences ^31^. In the presence of 1 mM MgCl_2,_ PAR_15_ underwent a 24% increase in base stacking events, whereas PAR_22_ exhibited a 103% increase (Fig. 4E). This greater frequency of π-π interactions between adenine bases could contribute to the greater compaction of PAR_22_ compared to PAR_15_.

### PAR_22_ Displays ADP-ribose Bundles

To further analyze the structural ensembles revealed by EOM, we parameterized the structures according to their {R_g_,R_EE_} values and performed hierarchical clustering ^39^. Using this approach, we found that the ensembles are highly heterogeneous: R_g_ and R_EE_ values covered 30 Å and 100 Å ranges, respectively. Through hierarchical clustering, we grouped structures into clusters with similar size, ranging from highly extended to highly compact (Fig. S7).

To elucidate unique conformational features within these clusters, we computed heatmaps of pairwise distances between bases in EOM-selected structure ensembles (Fig. S8). These heatmaps revealed how ADP-ribose bases are connected along the PAR chains. In these maps, off-diagonal regions with shorter distance implies the crowding of distal bases. For PAR_15,_ the heatmap revealed proximity mainly along the diagonal of the heatmap. This trend progressed monotonically towards the corners, suggesting that the molecule predominantly adopts relatively featureless extended conformations, irrespective of the presence or absence of MgCl_2_ (Fig. S8A). In contrast, PAR_22_’s heatmap showed significant connections, or close contacts, between bases that are close together (blue regions, slightly off-diagonal). One such region appeared in the 100 mM NaCl map (upper left corner), while two were evident when 1 mM MgCl_2_ was added (upper left and lower right corners, Fig. S8B). These off-diagonal regions indicate a local bundle of non-adjacent bases at the end(s) of the molecule, corroborating findings of local compaction initially identified in our MD simulations (Fig. 2).

We next considered the role of π-π stacking in these distinct ensembles. For PAR_15_, regions of proximity (*i*.*e*., low inter-base distance) correlated somewhat with where base stacking occurs, mainly along the diagonals (Fig. S8A). Yet, for PAR_22_, these off-diagonal low-distance regions were not enriched with base stacks (Fig. S8B). Most stacking events occurred between adjacent bases along the chain. Thus, while PAR_22_ has more base stacking with 1 mM Mg^2+^ than PAR_15_ in general (Fig. 4E), the observed ADP-ribose bundles appear unrelated to base stacking. Rather, local chain compaction due to the proximity of the PAR phosphate backbone to an Mg^2+^ ion likely triggers the intra-chain coil-to-globule transitions that lead to these bundles. This transition is evident in individual frames of the MD simulations, where the compaction correlates to some extent with proximity of Mg ions to the backbone (Fig. S9, Movie S2). Taken together, our analyses on the heterogeneous structural ensembles confirm the presence of ADP-ribose bundle formation unique to PAR_22_.

### Distinct Backbone Conformations for PAR_15_ and PAR_22_

Hierarchical clustering partitioned the structural ensembles into groups based on the overall size of each structure; however, deriving a more concise description of the PAR backbone conformations was challenging due to a variety of poorly related structures populating in any size subgroup (Fig. S7). Inspired by 2D classification of structures in single particle cryo-electron microscopy ^40^, and graph theory, we grouped the ensembles into unique, interrelated conformational subclasses. By applying 3D spatial alignment into network graphs and performing spectral clustering, we captured all conformations present in the ensembles, with no graph outliers (Fig. S10). The low spatial variation between the constituent structures of these subclasses (Fig. 5 and S11) supports that our algorithm effectively identified sensible classes, justifying the subsequent averaging to depict a single representative conformation in each class.

**Figure 5.**
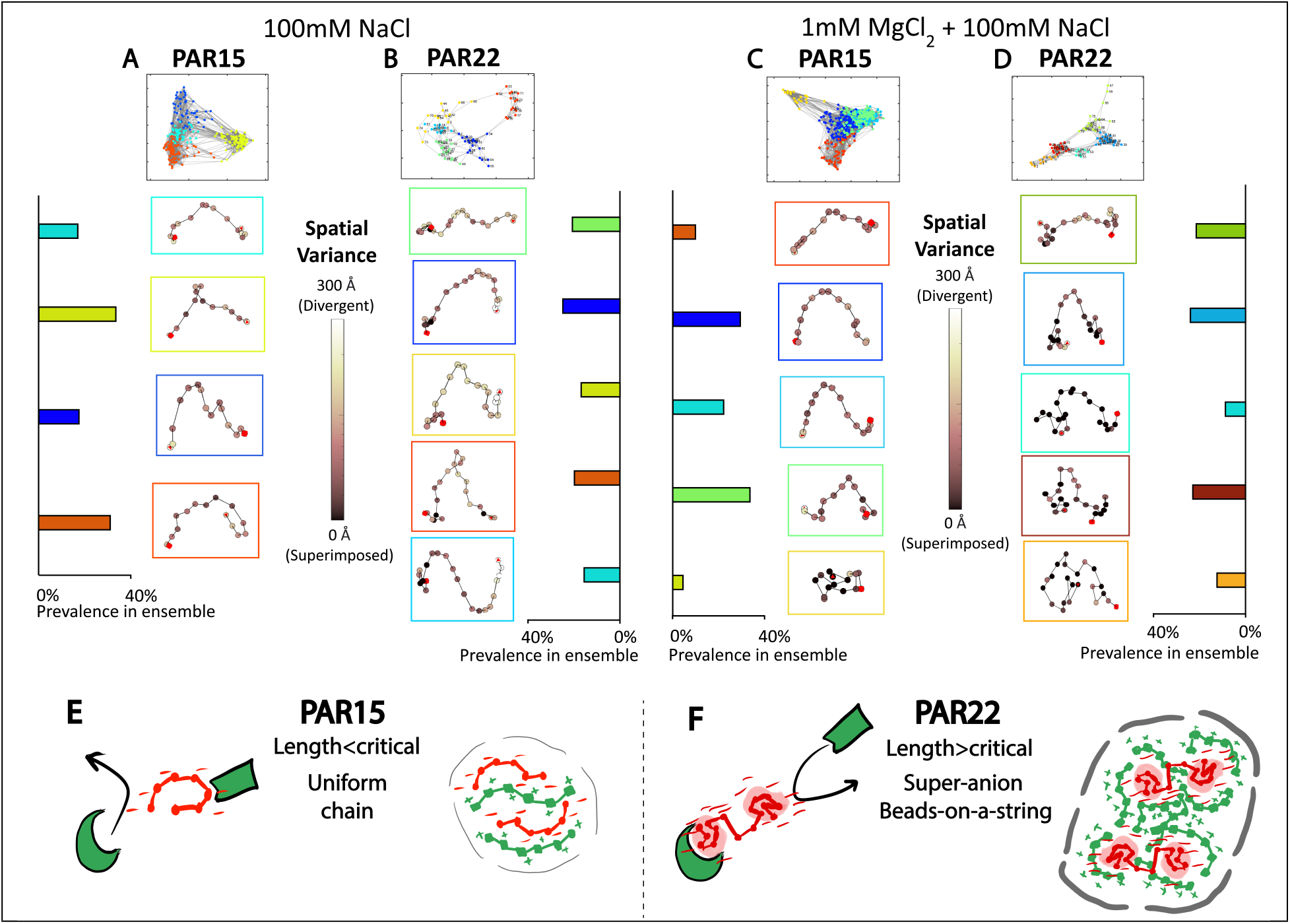
Backbone structural features of PAR_15_ vs PAR_22_ identified through spectral clustering. **(A-D)** PAR_15_ and PAR_22_ in 100 mM NaCl, both without and with the presence of 1 mM MgCl_2_ is shown Top plots show the graphs of PAR structures in each ensemble, color coded by the clusters identified by K-means. The box colors around each identified subclass matches their locations in the graphs. Wireframe models show the mean PAR backbone conformation in each case – each dot represents the mean position of each pair of phosphorus atoms across the entire subclass, colored by the degree of spatial variance present across that class. Red squares denote the 1” ends of the backbones and red triangles denote the 2 ʹ ends. The fraction of each structural subclass within the entire ensemble is shown adjacent to the respective averaged backbone conformer models. **(E-F)** Proposed model of a critical length for coil-to-globule transitions in PAR in the presence of MgCl_2_, linking to previously observed differences in molecular recognition of ligands and condensate formation in complex with positively charged species ^16^.

Without MgCl_2_ in 100 mM NaCl, the subclasses identified for PAR_15_ largely displayed similar conformations, with the backbone predominantly bent slightly into an inverted U shape (Fig. 5A). PAR_22_ exhibited similar U-shaped bends, but with additional variations: 21.3% of its conformations were more extended (Fig. 5B, green) and 16.0% more twisted (turquoise), likely contributing to the observed increase in tortuosity (Fig. 4E). Notably, bundles of ADP-ribose units were observed exclusively in PAR_22_ at the 1” ends of each subclass (Fig. 5B), visually confirming our pairwise distance measurements between bases (Fig. S8).

The introduction of 1 mM MgCl_2_ accentuated the conformational differences between PAR_15_ and PAR_22_. The structural ensembles at 100 mM NaCl of both PARs were generally less connected, exhibiting greater distances between pairs of structures (Fig. S10A-B). However, the presence of Mg^2+^ led to greater similarity among the structures within the ensemble, as evidenced by the increased number of structures demonstrating low root mean square deviations in pairwise comparisons (Fig. S10C-D). Specifically, in PAR_22_, the occurrence of ADP-ribose bundles was now noted in all five of the identified subclasses, spanning those with extended and more compact conformations (Fig. 5D). This observation is consistent with the heatmap indicating an increase in the number of regions that have short pairwise distance between bases (Fig. S8). The consistent low spatial variance (< 20 Å) in these bundle regions across all subclasses further indicates the systematic presence of bundles throughout the structural ensemble (Fig. 5D). Each bundle contained approximately eight ADP-ribose units at each end, interconnected by six additional units.

In contrast, such bundles were not observed in PAR_15_ with 1 mM MgCl_2_. These conformations closely resembled PAR_15_ in 100 mM NaCl alone, exhibiting similar backbone bending. A small fraction (4.6%) of the structures collapsed into a globule (Fig. 5C, yellow), akin to the most compact conformation observed in our initial MD simulations (Fig. 2). This subclass of collapsed globules may account for the slight 5.3% decrease in R_g_ in PAR_15_ as observed through SAXS (Fig. 3B). These data imply that only a small subset of molecules could undergo relatively featureless collapse with the small amount of Mg^2+^ present, leaving the rest of the ensemble relatively uninfluenced. On the other hand, the widespread bundling of the ADP-ribose units in PAR_22_ could explain the larger 19.0% decrease in its R_g_ (Fig. 3B). The difference in ADP-ribose bundle appearance alludes to a model of PAR’s length-dependent function (Fig. 5E,F).

### PAR Has Less Helicity and Base Stacking Than Poly-adenosine RNA

Our characterization of PAR and its distinct structural features prompted us to draw comparison with poly-adenosine RNA. Though composed of the same ribose, phosphate, and base building blocks, poly-A RNA and PAR have vastly different cellular functions. The former largely acts as a termination signal and binding motif, while the latter functions as a flexible binding scaffold. To delve into structural difference between these two nucleic acids likely tied to their divergent functions, we compared a 15-mer of ADP-ribose (PAR_15_) to a 30-mer of AMP (rA_30_) RNA. Both molecules were measured with SAXS in identical solutions containing 100 mM NaCl. Because ADP-ribose (in PAR) contains twice the number of phosphate and ribose groups as AMP (in RNA), these two macromolecules have comparable length and overall charge. Their R_g_ values further affirmed their similarity (Fig. 6A). Importantly, we chose PAR_15_ for comparison due to its lack of ADP-ribose bundles (Fig. 5), a feature not known to be present in poly-A RNA.

**Figure 6.**
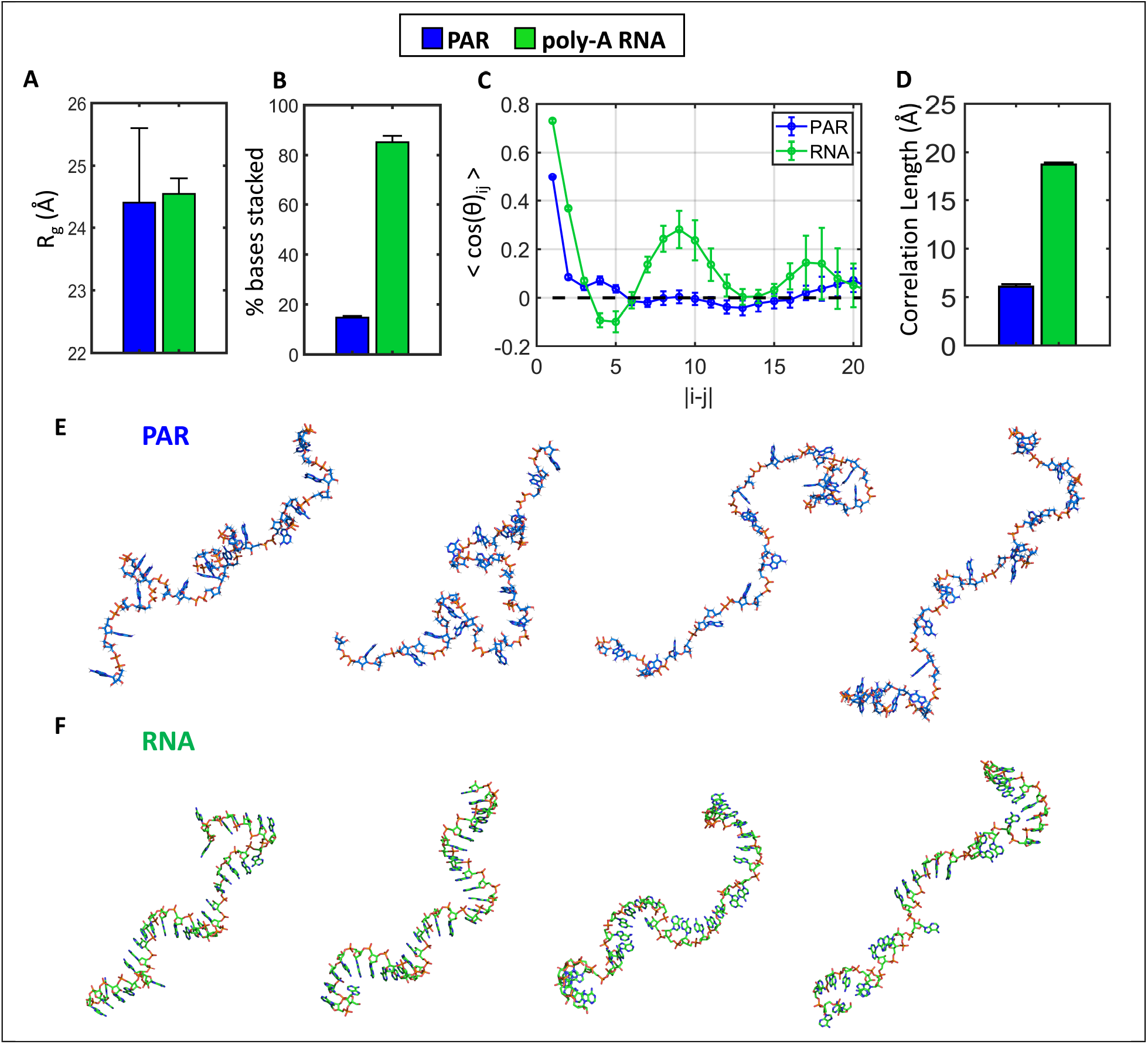
Polymeric differences of PAR_15_ vs poly-adenine RNA (rA_30_) in 100 mM NaCl. **(A)** SAXS-derived R_g_ of PAR_15_ vs rA_30_. **(B)** Fraction of adenine bases that are stacked in the PAR_15_ and rA_30_ structural ensembles. **(C)** Ensemble-averaged orientation correlation functions of PAR_15_, compared to that of rA_30_. Error bars represent the variance in the mean. **(D)** Correlation lengths of PAR_15_ vs rA_30_, computed across the structural ensembles. **(E)** Four structures from the conformational ensemble of PAR_15_ that are most highly selected by EOM. **(F)** Four representative rA_30_ structures; accessible via SASDFB9 in the Small Angle Scattering Biological Data Bank ^31^.

The orientational correlation function (OCF) can describe the orientational alignment of local regions of a polymer chain as a function of distance between its monomers (|i-j|). Peaks in OCF signifies high periodic orientational directionality, while a featureless exponential decay represents random chain orientations ^41^. Previous structural characterization of rA_30_ has revealed its well-ordered helix form, attributed to the propensity of adenine bases to undergo π-π interactions. The helix formation is driven by an extensive base stacking network, with 85.2 ± 2.5% of the bases adopting a parallel stacked configuration (Fig. 6B) ^31^. The OCF of rA_30_ displayed strong oscillatory behavior with peaks spaced by the periodicity of an A-form helix (Fig. 6C). In contrast, PAR_15_ exhibited less orientational correlation along the chain, with the exponential decay of its OCF more closely resembled that of a random coil at |i-j| > 4 (Fig. 6C) ^41^. This difference suggests that the bases in PAR are more randomly arranged than in RNA, supported by the mean PAR correlation length (6.1 ± 0.3 Å) being only a third of that of poly-A RNA (18.7 ± 0.2 Å) (Fig. 6D). The correlation length, which is greater when repeating backbone orientations are present, reflects the lower degree of order for PAR. Moreover, less than 14.4 ± 0.9% of the bases in PAR_15_ were stacked (Fig. 6B), preventing π-π interactions from stabilizing an ordered helical conformation, as observed in rA_30_. The additional phosphate group and ribose sugar between each adenine base in PAR, compared to poly-A, may place the bases too far apart for extensive π-π base stacking interactions. Such differences in helicity and base stacking were evidently observed in individual sample conformers (Fig. 6E-F).

## Discussion

The building blocks of PAR are configured uniquely compared to other nucleic acids, potentially contributing to its distinct functional properties. Previously, we have shown that PAR possesses a larger persistence length than RNA, thereby being stiffer—a characteristic attributed to the different distribution of the phosphates in PAR relative to RNA ^30^. Both PAR and RNA assume more compact states with increasing salt concentration—a compaction that occurs with 100x fewer divalent than monovalent ions ^30^. Building on this polymeric characterization, we set out to explore the 3D structure of PAR. Our studies reveal that the structure of PAR markedly diverges from that of poly-A RNA. In contrast to RNA, PAR possesses a shorter correlation length, exhibits less π-π stacking between bases, and adopts a less helical structure (Fig. 6). These findings are aligned with previous studies showing that PAR, when exposed to only monovalent ions at room temperature, lacks a well-defined secondary structure ^17–21^ While the two delocalized rings in adenine make them especially prone to undergo a π-π interaction, the enhanced electrostatic repulsion in PAR, relative to RNA, could also prevent bases from achieving the close proximity needed to form such an interaction. This distinction may align with PAR’s unique ability, compared to RNA, to bind positive ligands, both in 1:1 interactions and large-scale condensates ^3,11,12,16^.

PAR has historically presented challenges for structural characterization. The few published studies on PAR structure suggested that it lacks a well-defined structure but may have some subtle structural features ^17,21,22^ However, a common limitation among these studies is their reliance on data derived from mixtures of PAR lengths, potentially masking signals from individual lengths. In this study, we have synthesized homogenous, single-length PARs for structural investigation. We have integrated structures generated through PAR_15_ and PAR_22_ MD simulations with experimental data offered by SAXS. With this approach, we can ascertain and analyze conceivable heterogenous ensembles of structures, aiming to comprehend not just how PAR interacts with the surrounding ions but also the impact of length on its structural heterogeneity.

SAXS data revealed that PAR_22_ compacts more than PAR_15_ with Mg^2+^ (Fig. 3). Analysis of order parameters in the SAXS/MD-derived structural ensembles further elucidates differences in Mg^2+^-induced PAR compaction between 15- and 22-mer. The additional compaction observed in the latter is driven by a more tortuous backbone (Fig. 4D) and increased stacking among adenine bases (Fig. 4E). These phenomena are mediated by Mg^2+^-induced transient contacts with the PAR phosphate backbone, causing local distortion and leading to the more compact configuration observed in the MD trajectories (Fig. S9, Movie S1 and S2).

By examining subclasses of the full structural ensembles, we characterized the presence of ADP-ribose bundles unique to PAR_22_ (Fig. 5). This phenomenon aligns with a prior MD study that observed multiglobular behavior for a 25-mer, based on the dihedral constraints of the molecule ^22^. However, the previous study, using improper force fields without corrections for ions, resulted in more compact PAR structures than realistically expected. Our 1 µs-simulations demonstrate that such compact states can be long-lasting (50-100 ns), oscillating transiently between compact and extended states (Fig. S9). This behavior, especially under conditions with divalent cations or at higher monovalent cation concentrations, was not observed in the previous study due to its methodological limitations. Specifically, the heightened complexity inherent in simulations featuring divalent cations, like Mg^2+^ ions, was not addressed, underscoring the significance of our study in revealing the dynamic nature of PAR structures.

Our approach, combining experimental data with MD simulation, reveals that these ADP-ribose bundles are seeded by the presence of Mg^2+^ (Fig. S9), and these features are less pronounced in PAR_15_. Once formed, the structural stability of these bundles does not seem to rely on adenine base stacking or direct association with divalent Mg^2+^ ions alone. Instead, Na^+^ ions within the bundle appear to play a stabilizing role (Movie S2). We speculate that a minimum number of ADP-ribose units are required to form a stable bundle—a critical length not attained by PAR_15_—possibly due to insufficient length between bundles to adequately separate congregated negative charges. Ensemble-level pairwise base distance heatmaps support the observation that globular bundles of ADP-ribose units are diffusely present across the entire ensemble in PAR_22_, but not in PAR_15_ (Fig. S8). It is conceivable that in longer lengths of PAR (>50mer), multiglobular ADP-ribose bundles could periodically appear along the chain, consistent with the beads-on-a-string models of classical polymer theory ^42,43^.

Based on these findings, we propose a model that addresses unresolved questions concerning PAR’s diverse interactions with its binding partners. The model suggests the formation of globular ADP-ribose bundles beyond a crucial length, which falls between 15 and 22 units. These bundles appear to form through an intra-molecular coil-to-globule phase transition induced by divalent cations, wherein PAR of sufficient length can have part of its chain as a collapsed globule and the rest as an extended polyelectrolyte. This phenomenon was characterized through our MD simulations, a detail that would have been overlooked by simply observing the average system behavior. At physiological cationic conditions, we observe that PAR undergoes this transition rapidly on the sub-microsecond timescale. We speculate that these bundles could be of variable size, thereby granting PAR more conformational variety in molecular recognition of binding partners (Fig. 5E-F) ^17^. Additionally, these bundles localize the negative charge within the molecule, forming super-anion beads-on-a-string in a length-dependent manner which could afford enhanced condensate formation (Fig. 5E-F) ^16^. Follow-up studies are warranted to test these hypotheses.

### Limitations

Our focus has been on characterizing linear PAR—a choice driven by the current capability in the field to synthesize sufficient, pure quantities of this single defined length molecule, essential for interpretation of SAXS experiments. Exploring branched PAR, another physiological structural form, is not technically feasible at this juncture. Moreover, synthesizing and simulating specific lengths of long PAR—potentially reaching up to 200-mer in cells—poses notable challenges. Under physiological conditions, PAR is primarily covalently conjugated to proteins, rather than existing as freely diffusing molecules. Given that proteins are conjugated to the ends of PAR, we postulate that ADP-ribose bundles might form in the middle of a chain, possibly exhibiting a periodicity of 15-22 units, rather than exclusively at the ends. Future systematic studies on PAR of different lengths, structures, and conjugations to proteins are essential for a more comprehensive understanding of the structure and function of PAR.

## Conclusions

In this study we characterized the structure of PAR_15_ and PAR_22_ by performing MD in conjunction with SAXS and carrying out detailed analyses on the resulting conformational ensembles. We showed that PAR rapidly compacts with increasing ionic strength, and that this compaction occurs differently in the two different PAR lengths. The structural ensembles of PAR were found to be highly heterogeneous. By breaking down this heterogeneity through biophysical parameters and real space class averaging, we characterized the conformational mechanism for this difference in compaction, identifying globular bundles of ADP-ribose unique to PAR_22_. We speculate that this structural feature may enable PAR to bind ligands specifically, forming part of the PAR code ^44^.

## Materials and Methods

### Molecular dynamics (MD) simulations

All MD simulations were performed using NAMD2.14 ^45^, the CHARMM36 parameter set for protein and DNA ^46^, TIP3P water model ^47^, and a custom hexahydrate model for magnesium ions along with the CUFIX corrections to ion–nucleic acid interactions ^23^. Multiple time stepping was used: local interactions were computed every 2 fs, whereas long-range interactions were computed every 6 fs ^48^. All short-range nonbonded interactions were cut off starting at 1 nm and completely cut off by 1.2 nm. Long-range electrostatic interactions were evaluated using the particle-mesh Ewald method computed over a 0.11 nm spaced grid ^49^. SETTLE and RATTLE82 algorithms were applied to constrain covalent bonds to hydrogen in water and in non-water molecules, respectively ^50,51^. The temperature was maintained at 300 K using a Langevin thermostat with a damping constant of 0.5 ps^-1^, unless specified otherwise. Constant pressure simulations employed a Nose-Hoover Langevin piston with a period and decay of 200 and 50 fs, respectively ^52^. Energy minimization was carried out using the conjugate gradients method. Atomic coordinates were recorded every 9.6 picoseconds, unless specified otherwise. An example MD timeseries plot is shown in Fig. 2B. Visualization and analysis were performed using VMD and MDanalysis ^53–55^.

CHARMM-compatible force field parameters for PAR were obtained by combing existing parameters for chemically similar moieties. Specifically, a custom patch was written to define a PAR residue (PAR), whereby the oxygen atom on the terminal phosphate group of an ADP molecule (atom O5D) was connected to the ribose sugar (atom C5D) using analogy from NADP. 0 patch (BND) connected the ribose (atom O1D) with ADP (C2ʹ). With these two patches, a PAR molecule of arbitrary number of monomers could be defined. Separate patches were defined for the terminal atoms, a hydrogen on the O1D atom of the ribose (1TER) and an OH group on the C2ʹ atom of ADP (2TER). The topology and parameter files for a PAR residue are provided in Supplementary Materials.

### Small-angle X-ray scattering (SAXS)

Solution SAXS experiments on PAR_15_ and PAR_22_ were performed at the ID7A1 BioSAXS beamline at the Cornell High Energy Synchrotron Source and the 16-ID Life Science X-ray Scattering beamline at the National Synchrotron Light Source II of Brookhaven National Laboratory ^56,57^. Radial integration and data reduction was performed in BioXTAS RAW to obtain 1D scattering profiles, plotting scattering intensities (I, in arbitrary units) as a function of momentum transfer (q, in Å^-1^) ^58^. PAR was assayed at 60µM, 40µM, and 20µM concentrations to observe for interparticle effects at low q, and these were corrected when present by linearly extrapolating each point at q < 0.05 Å^-1^ to the zero-concentration limit. Radii of gyration were obtained through Guinier analysis ^41^, approximating the scattering profile at (q*R_g_) < 1.3 as a Gaussian function:

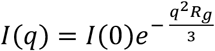

Guinier analyses are shown in Fig. S4. Data were plotted in the size-normalized Kratky format to emphasize changes in overall macromolecular shape. The molecular form factor (MFF) model was fit to the scattering data and used to describe the arrangement of the PAR chain ensembles relative to the theta state through the Flory (v) parameter (Fig. S3) ^34,59^. Scattering data are deposited on the Small Angle Scattering Biological Data Bank (SASBDB) with the identifiers SASDSJ5, SASDSK5, SASDSL5, SASDSM5.

### Determining PAR structural ensembles

To provide a diverse pool of structures for PAR, we sampled the MD trajectories in 1 nanosecond snapshots, obtaining a set of 1000-1200 individual structures over a broad, continuous conformational range. Fig. S5 shows the structural ensembles parameterized in Cartesian space by their R_g_ and R_EE_. CRYSOL v2.0 was used to compute the theoretical scattering profile of all structures in the SAXS regime, with a maximum order of harmonics of 15, Fibonacci grid order of 18, 0.3 Å^-1^ maximum scattering angle, and 61 calculated data points ^60^.

The raw MD simulations were seen to approximate well the overall size of the conformational ensembles but not their overall shape. This is evidenced by the discrepancies between the summed theoretical scattering of all structures in the pool and the experimental scattering beyond the low q regime (q > 0.03 Å^-1^) (Fig. S6). To provide an accurate depiction of the solution-state ensemble of structures that the molecules sample, the information content of the SAXS data was leveraged through EOM v2.1 (ATSAS, EMBL Hamburg, Germany) ^37,38^. Briefly, subsets of the pool of structures were sampled and the summated theoretical SAXS profile of the subset of structures was computed and fit to the experimental SAXS profile – the process was iterated until agreement with the experimental data is reached. EOM was run over 1000 generations with 50 ensembles, 20 curves and 10 mutations per ensemble, over 100 iterations. This was found to yield convergence of the fit to the experimental SAXS data, with χ^2^ values of 0.1 – 0.4 and the SAXS profiles of the final pool post-EOM falling within error of the experimental measurement across the entire q range (Fig. S6). Introducing more iterations into the algorithm beyond that executed was not found to improve convergence of the χ^2^ meaningfully, except for PAR_22_ in 100 mM NaCl where 10000 iterations were employed. The final ensembles were assessed to fall along a continuous distribution in their R_g_, R_EE_, and maximal dimensions, with no implausible bimodalities (Fig. S5).

## Acknowledgments

We thank Drs. Shirish Chodankar, Richard Gillilan, Qingqiu Huang, and Suzette Pabit for their invaluable support with SAXS data collection.

## Funding

This work is based on research conducted at the Center for High-Energy X-ray Sciences (CHEXS), which is supported by the National Science Foundation (BIO, ENG and MPS Directorates) under award DMR-1829070, and the Macromolecular Diffraction at CHESS (MacCHESS) facility, which is supported by award 1-P30-GM124166-01A1 from the National Institute of General Medical Sciences (NIGMS), National Institutes of Health (NIH), and by New York State’s Empire State Development Corporation (NYSTAR). The LiX beamline is part of the Center for BioMolecular Structure (CBMS), which is primarily supported by NIGMS, NIH through a P30 Grant (P30GM133893), and by the DOE Office of Biological and Environmental Research (KP1605010). LiX also received additional support from NIH Grant S10 OD012331. As part of NSLS-II, a national user facility at Brookhaven National Laboratory, work performed at the CBMS is supported in part by the U.S. Department of Energy, Office of Science, Office of Basic Energy Sciences Program under contract number DE-SC0012704. A.A. and K.C. acknowledge support from the NIH via R01-GM137015. The supercomputer time was provided by ACCESS allocation MCA05S028 and Leadership Resource Allocation MCB20012 on Frontera at the Texas Advanced Computing Center. Frontera is made possible by National Science Foundation award OAC-1818253. This work at the Leung lab is supported by the NIH grants T32-CA009110 (M.B.) and R01-GM104135 (A.K.L.L.). Work in the Pollack lab is supported by NIH grant R35-GM122514. T.W. is supported by a Natural Sciences and Engineering Research Council of Canada postgraduate scholarship (NSERC PGS-D).

## Author Contributions

Conceptualization: AKLL, LP, AA

Formal Analysis: TW, KC

Investigation: TW, KC

Resources: MB

Writing—Original Draft: AKLL, LP, TW

Writing—Review & Editing: AKLL, LP, AA, TW, KC, MB

Supervision: AKLL, LP, AA

Funding Acquisition: AKLL, LP, AA

## Competing interests

The authors declare that they have no competing interests.

## Data and materials availability

All data needed to evaluate the conclusions in the paper are present in the paper and/or the Supplementary Materials. Raw data for all scattering profiles of PAR and poly-A RNA are available in the SASBDB as indicated. Files utilized for MD simulations are provided in the supplementary materials. Computer scripts used for data analysis are available upon request.

## Notes

### Competing Interest Statement

The authors have declared no competing interest.

### Summary of Updates

The manuscript has underwent significant rearrangement to better streamline it. As well, the final figure has been removed and re-integrated with figure 5.

