## Supplemental Information for "Length-dependent Intramolecular Coil-to-Globule Transition in Poly(ADP-ribose) Induced by Cations"

**Supplementary Materials**

Supplementary Materials and Methods

Figs. S1 to S11

References S1 to S9

Mp4 files of MD trajectories of $PAR_{15}$ and $PAR_{22}$ in 100 mM NaCl with and without 1 mM $MgCl_2$:

Movie S1 – $PAR_{15}$ with $Mg^{2+}$

Movie S2 – $PAR_{22}$ with $Mg^{2+}$

Movie S3 – $PAR_{15}$ no $Mg^{2+}$

Movie S4 – $PAR_{22}$ no $Mg^{2+}$

Topology and parameter files for a PAR residue, used in MD simulations

### Supplementary Materials for

#### Length-dependent Intramolecular Coil-to-Globule Transition in Poly(ADP-ribose) Induced by Cations

Wang, Coshic, Badiee *et al.*

**This PDF file includes:**

Supplementary Text
Figs. S1 to S11

**Other Supplementary Materials for this manuscript include the following:**

Movies S1 to S4

#### PAR sample preparation

Monodisperse samples of $PAR_{15}$ and $PAR_{22}$ were produced as detailed in Badiee et al [30]. Briefly, PAR was produced as a nonspecific, multi-length mixture through a reaction involving the PARP5a catalytic domain onto histones followed by isolation of PAR molecules. Specific lengths of PAR were isolated from the multi-length mixture through high performance liquid chromatography.

Prior to solution scattering experiments, PAR samples were concentrated and buffer exchanged into 100 mM NaCl, 20 mM Tris-HCl (pH 7.4) using 3k MWCO Amicon microcentrifuge spin columns (Millipore Sigma, St. Louis MO, USA), using six centrifugation steps of 14000 xg for 15 minutes, maintaining 4°C temperature. Samples were then annealed by heating to 90°C for 5 minutes and snap cooling at 4°C for 20 minutes, then stored on ice until SAXS data collection. For samples requiring $Mg^{2+}$, the final concentration was spiked to 1 mM $MgCl_2$ by mixing in a 1M $MgCl_2$ stock solution minutes prior to SAXS measurements.

#### Partitioning structural ensembles into size categories

To break down the structural heterogeneity within the conformationally broad ensembles, we performed hierarchical clustering by computing all pairs of Euclidean distances between the structures in 2D $\{R_g, R_{EE}\}$ space, generating a dendrogram using '*cluster'* in MATLAB, and selecting a distance cutoff for group differentiation. The dendrogram distance cutoff was selected such that 3-4 unique clusters, each containing at least 10-40% of all structures in the ensemble, were yielded. These were further defined as low $R_g$, middling $R_g$, and high $R_g$ families of structures (Fig. S7).

Differences between families of $PAR_{15}$ and $PAR_{22}$ structures were determined by visualizing maps of pairwise distances between all bases. To do this, the base positions were sampled as the average coordinates of adenine base atoms in the PDB files and the Euclidean distance between all pairs of base positions were computed and plotted in a heatmap as a function of base number along the chain.

#### Structural order parameters to characterize PAR structures

To further describe the structural features of the determined PAR conformer ensembles and to compare them to the previously characterized behavior of poly-adenine RNA, we computed the tortuosities, number of base stacking events, and correlation lengths. These parameters were computed as weighted averages in each structural ensemble, with weights given by how many times each conformer was selected by the EOM algorithm. The coordinates of the average position of each pair of Phosphorus atoms and the Oxygen atom in between each pair of ribose sugars were sampled along the PAR chain to break it into even segments. Tortuosity indices (T) were computed as [61]:

$$T = \frac{\sum_{i=1}^{n-1} \theta_i}{L}$$

Where we take the sum of all n-1 bond vector angles $\Theta_i$ along the sampled chain, where n is the total number of ADP-ribose monomers, and divide by the end-to-end distance L. T was computed for all structures in the PAR ensembles and comparison of tortuosities were performed through a two sample T-test assuming unequal variances, with a p-value threshold of 0.05 considered significantly different.

The orientation correlation function (OCF) was employed to visualize structural features across the selected PAR conformer ensembles [41]. The OCF was calculated as:

$$OCF = \langle cos\theta_{ij} \rangle = \langle \hat{r}_i \cdot \hat{r}_j \rangle$$

Where r is the normalized bond vector between each pair of sampled coordinates. OCFs were plotted as a function of the distance between linkages |i-j|, for all {i,j} pairs. To get a metric of approximate polymer stiffness and how structured the chain is, correlation lengths ($l_{OCF}$) were computed by summing across the OCFs:

$$l_{OCF} = b \sum_{ij}^{n-1} OCF(i,j)$$

Where b is the length of bonds between sampled coordinates. OCF curves were truncated to |i-j|=20 for visualization – past this, greater OCF error made their comparison difficult.

Lastly, to determine the prevalence of π- π interactions in stabilizing the PAR chains, the number of base stacking events was computed for each structure. Adenine bases were indexed by sampling the coordinates of base atoms and fit using a plane. Pairs of base planes were considered stacked when the normal vectors to each plane were both < 5 Å apart and approximately collinear, with < 45° separation. These parameters were optimized such that they reproduced results from previously characterized structures of poly-A RNA that were derived from dinucleotide libraries with base stacking information built in.

All structural descriptors and order parameters were computed using in-house software written in MATLAB R2021a (Mathworks, Natick, MA, USA).

**3D classification and averaging of disordered PAR ensembles**

To break the multitudinous structural ensembles into a small subset of conformations, spectral analysis was performed to identify 4-5 unique conformational subclasses representative of the entire population. This allowed for visualization of the structural pool in real space in a practical manner. First, PAR structural ensembles were rotationally aligned using PyMOL v2.4.0 (Schrodinger, LLC) and the resulting root mean square deviation (RMSD) between each pair of aligned structures were input into an $N_{structures}$ x $N_{structures}$ matrix. Within this matrix, an RMSD threshold was set between 10-15 Angstroms, wherein a pair of structures were declared to be connected if the RMSD between them was below threshold. In this way, an adjacency matrix was constructed, which was input into the spectral clustering algorithm of Ng, Jordan, Weiss to map the structures to graph space wherein pairs of structures (nodes) are connected if they fall below the RMSD threshold [62]. Unique classes in the graph were then identified through K-means clustering in MATLAB, where it was found that 4-5 clusters sensibly categorized the node set. Upon classifying the PAR ensembles into classes, the structures were coarse grained by sampling the average positions of each pair of phosphorus atoms along the backbone, further aligned by RMSD minimization, and averaged to determine the characteristic 3D chain conformation of each subclass. Fig. S11 shows all structures overlayed with the average structure in each class, to showcase the validity of the method.

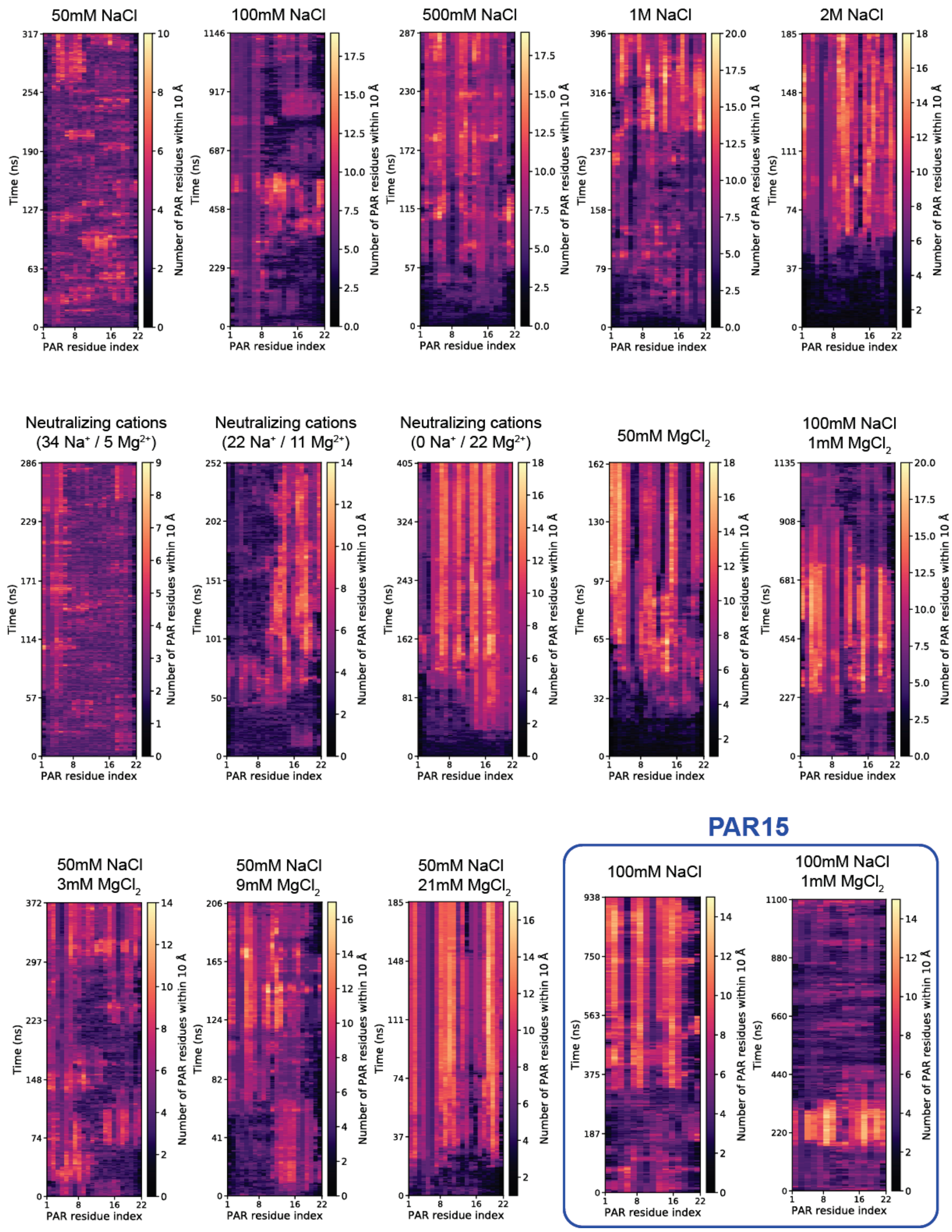


**Figure S1. Per residue contact analysis of MD trajectories.** For each simulation system, the heatmap displays the number of PAR monomers located within a 10 Å from the specified residue. The horizontal axis specifies the residue index of each PAR monomer, whereas the y-axis specifies the simulation time. The color of the voxels represent the number of contacts according to the color scale bar. All represented simulations are of $PAR_{22}$, except for the last two, which are of $PAR_{15}$

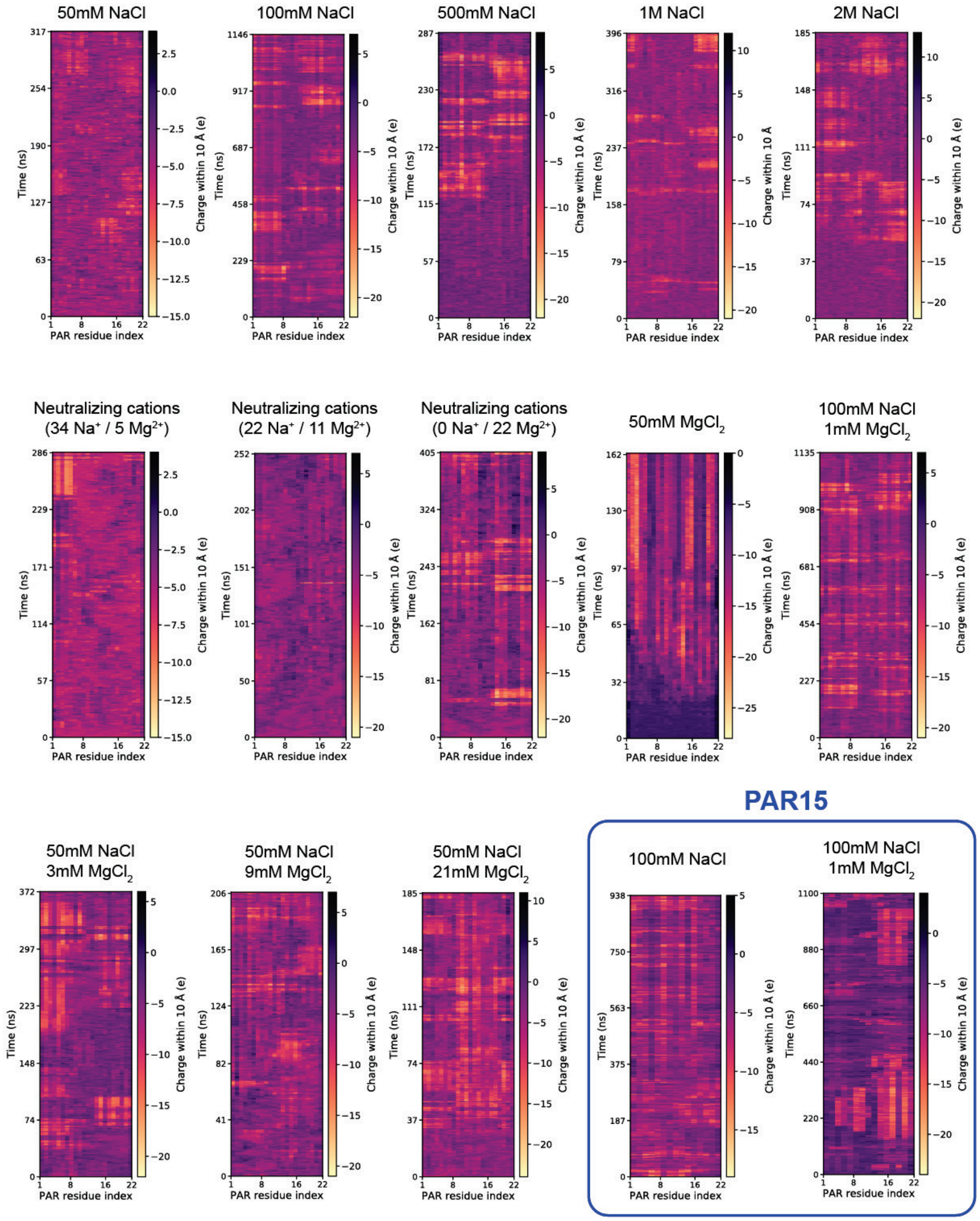


**Figure S2. Local compensation of PAR charge by counterion as seen in MD simulations.** For each simulation system, the heatmap displays the total electrical charge of all atoms located within 10 Å from the specified residue. The horizontal axis specifies the residue index of each PAR monomer, whereas the y-axis specifies the simulation time. Color of the voxels represent the local charge according to the color scale bar. All represented simulations are of $PAR_{22}$, except for the last two, which are of $PAR_{15}$.

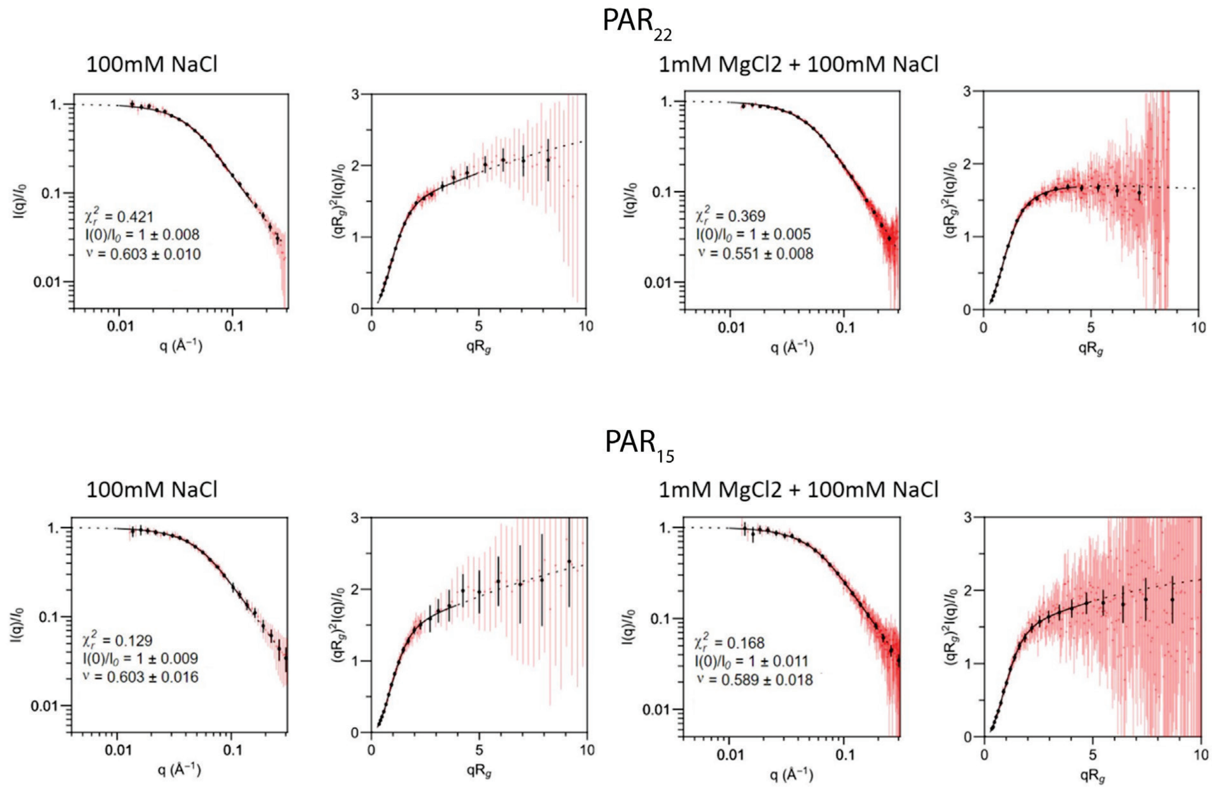


**Figure S3. Fits to the molecular form factor model to obtain ν.** Fits and plots were generated using the molecular form factor model and online resource [34].

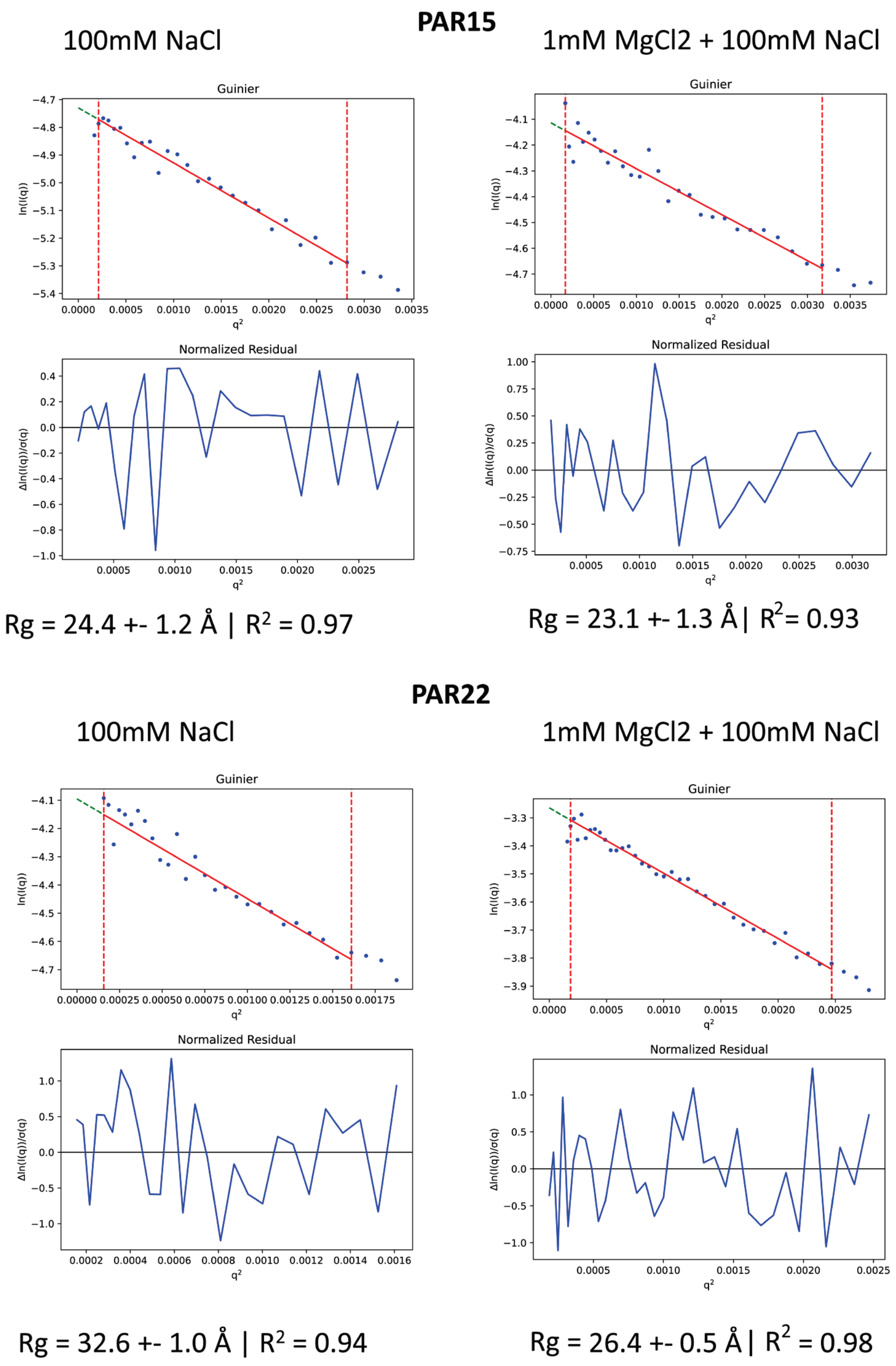


**Figure S4. Guinier fits[33] to determine $R_g$ of $PAR_{15}$ and $PAR_{22}$ in 100 mM NaCl with and without 1 mM $MgCl_2$.**

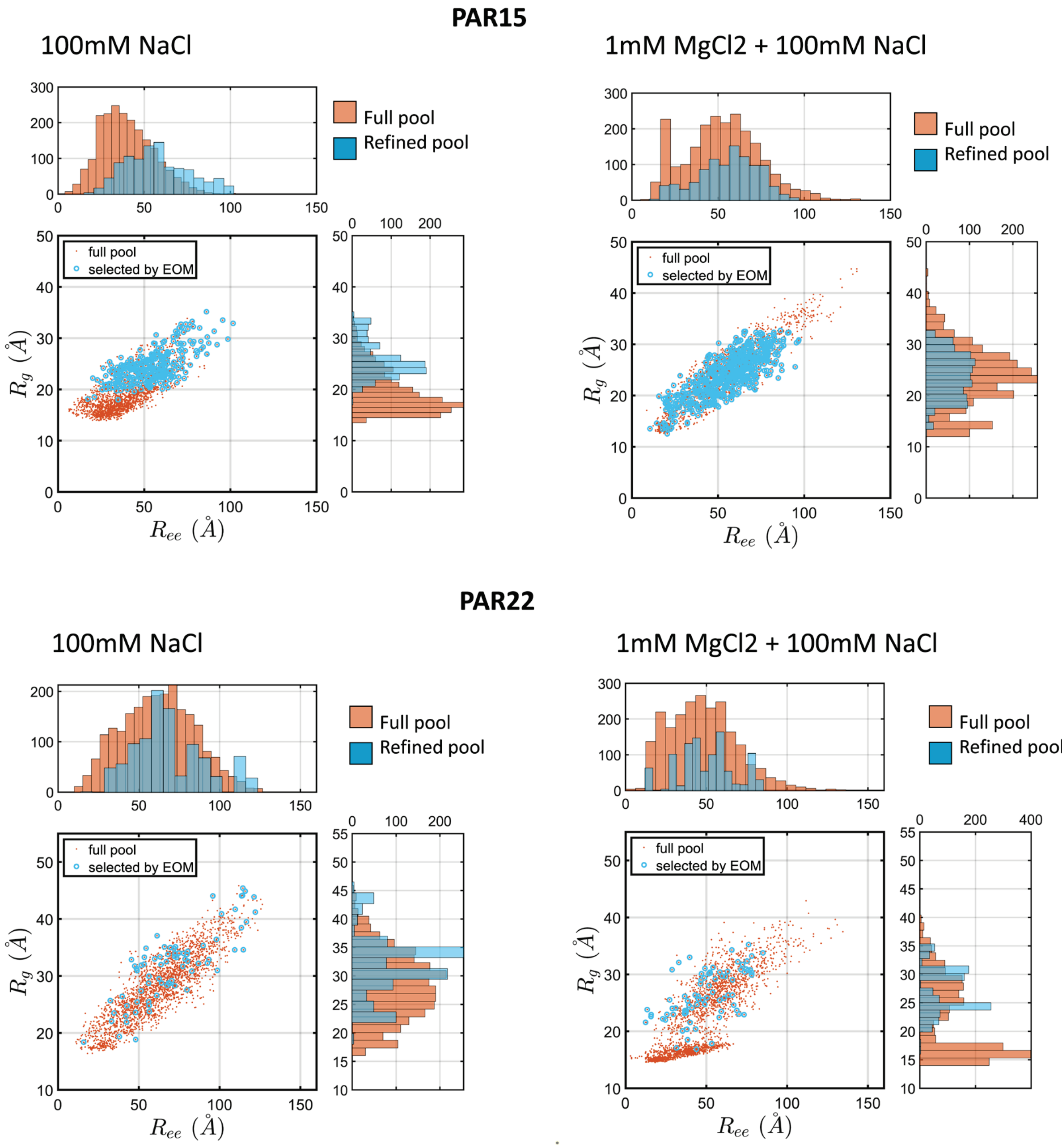


**Figure S5. Complete dataset for application of the ensemble optimization method[37,38] to determine structural ensembles for PAR.** The pool of structures in the full MD simulations are shown in orange and the ensembles determined using the SAXS data are shown in blue. Agreement to the experimental scattering are shown in Fig. S4.

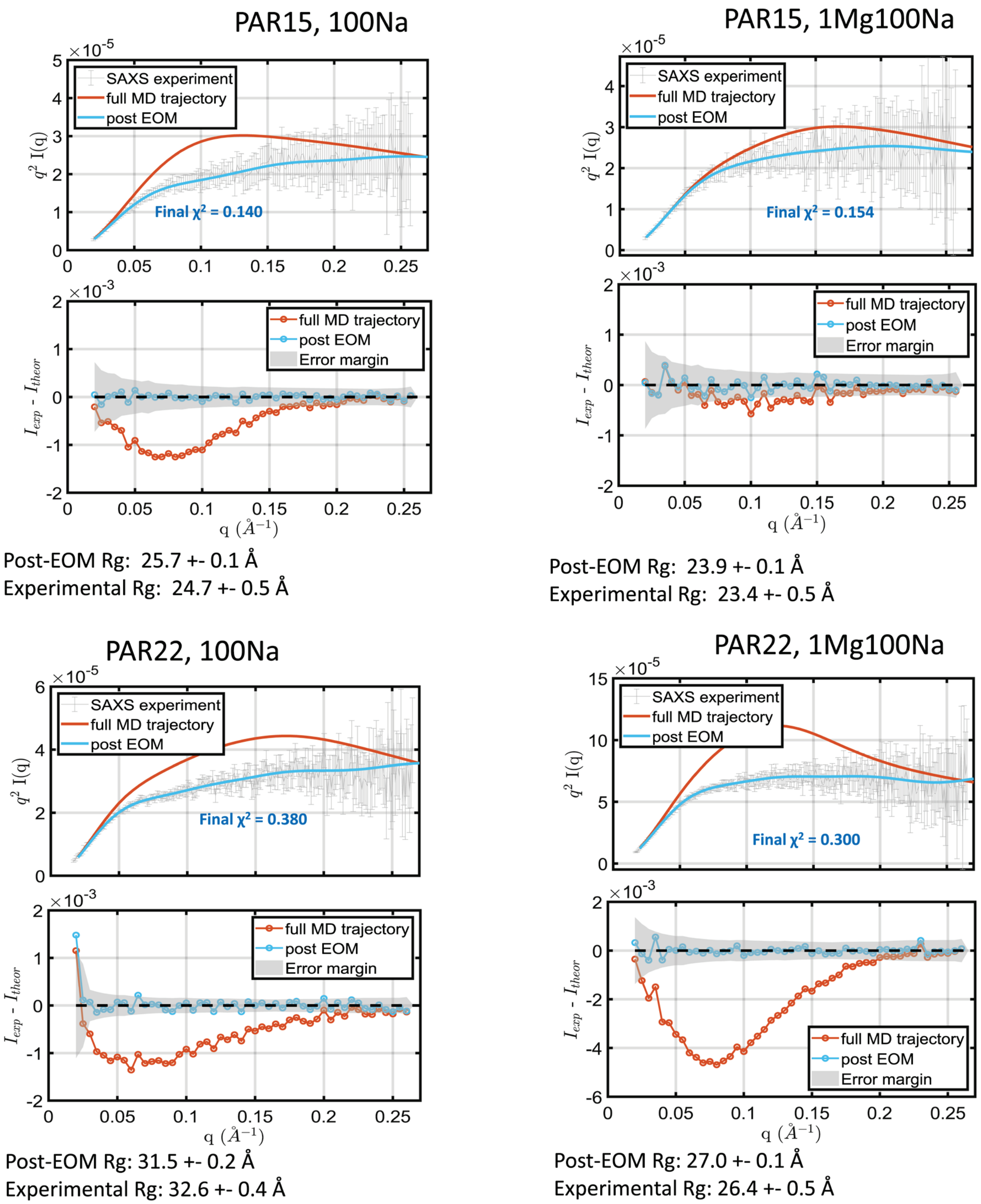


**Figure S6. Agreement to the SAXS data of the ensemble of structures in the full MD pool and the pool refined by EOM.** Agreement of theoretical MD and EOM pool scattering profiles with experimental data are shown in top plots on Kratky axes in gray with error bars. The bottom plots show residuals between theoretical and experimental scattering across the entire q range, as well as the experimental error margin in gray. The theoretical scattering post-EOM is shown to fall consistently within the experimental error margin. Mean $R_g$ values of the structural ensembles after EOM are also listed relative to the $R_g$ values determined from Guinier analysis.

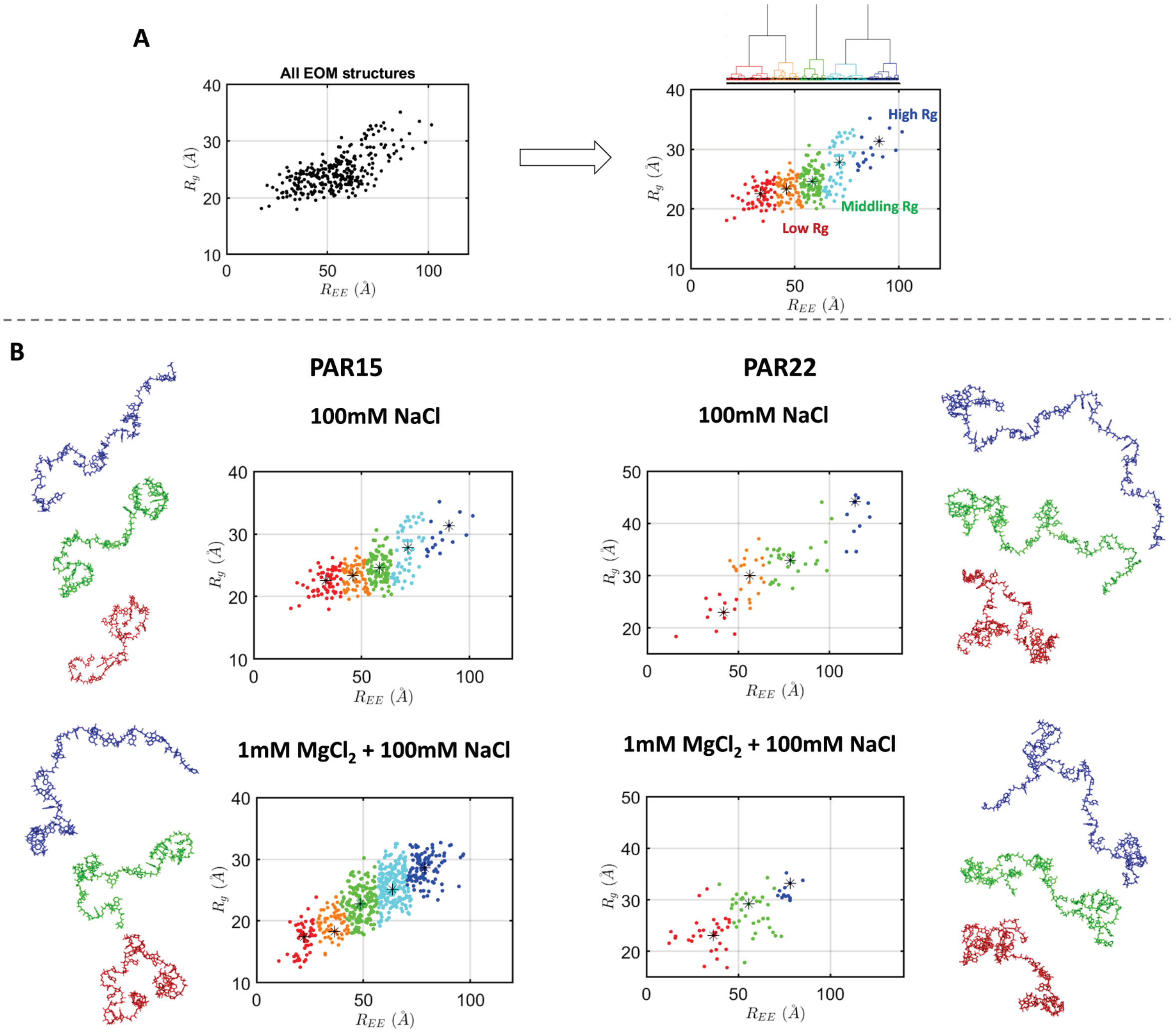


**Figure S7. Hierarchical clustering to identify unique subsets of structures within the full ensembles.** A) The general method of using dendrograms of Euclidian distances with the structural pool parameterized in 2D space. Representative structures are chosen as those closest to the centroid of each cluster., shown with a '*'. B) Structural groups for $PAR_{15}$ and $PAR_{22}$ in 100 mM NaCl, with and without 1 mM $MgCl_2$.

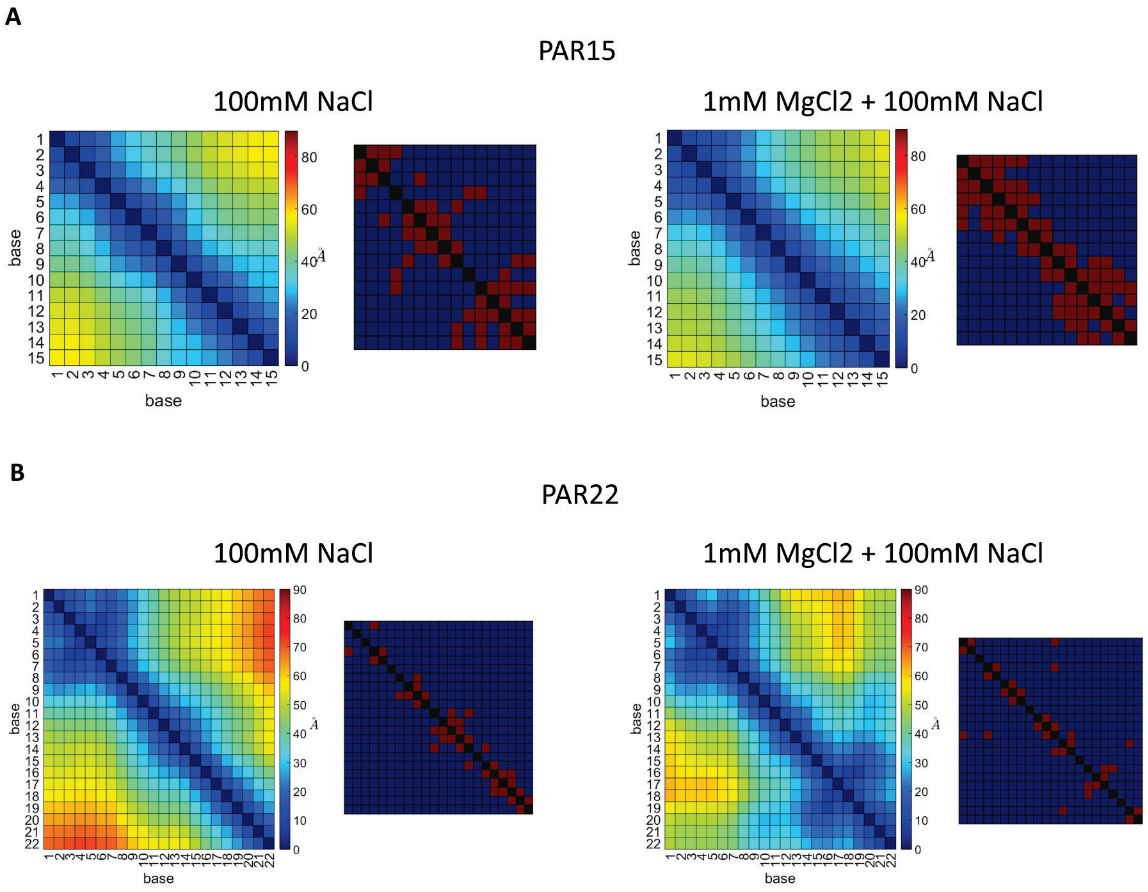


**Figure S8.** Heatmaps of average distances between all pairs of adenine bases across the PAR ensembles for A) PAR15 and B) PAR22, with and without $MgCl_2$. Shown to the right of each heatmap are binary base stacking matrices plotted on the same axes, showing which pairs of bases that base stacking is occurring. In these matrices, red represents a base stacking event between the corresponding pair of bases and blue represents that no base stacking is occurring.

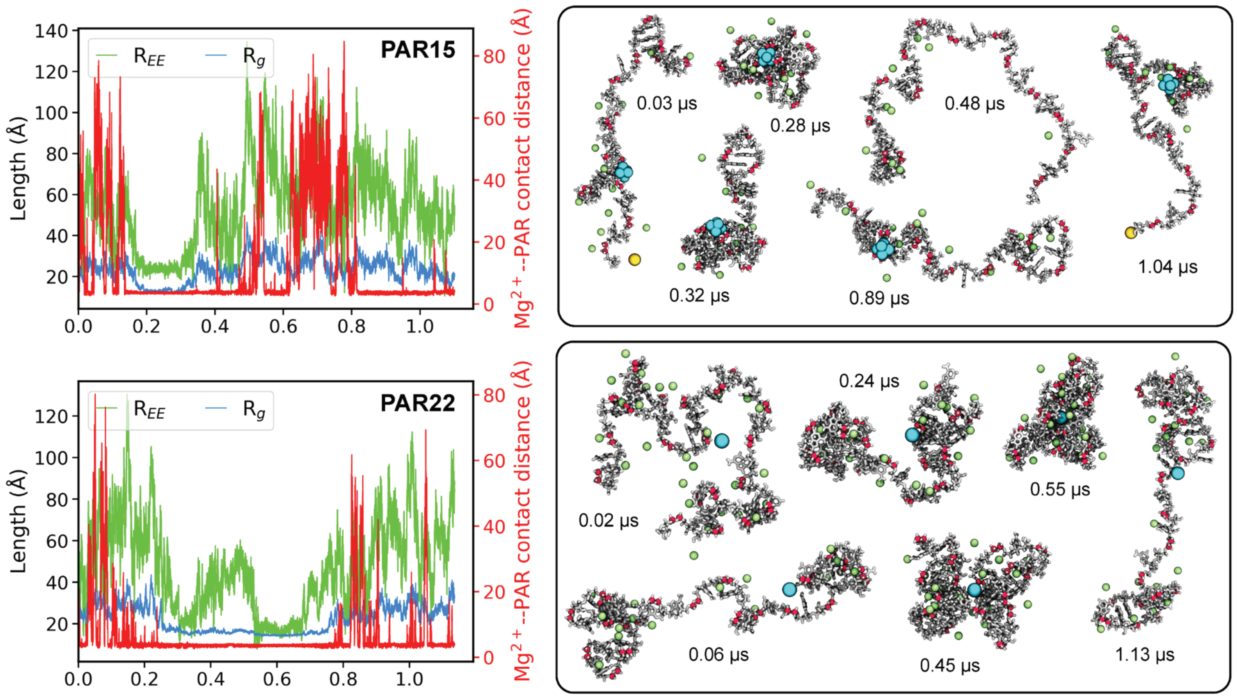


**Figure S9. $Mg^{2+}$ facilitates PAR compaction.** The minimal distance between a $Mg^{2+}$ ion and any non-hydrogen atom of PAR is recorded every frame and plotted as a function of the simulation time (red). The end-to-end distance ($R_{EE}$) and radius of gyration ($R_g$) are overlayed on the same plot. Top panels correspond to $PAR_{15}$ and lower panels for $PAR_{22}$. Snapshots from the simulation are shown in the enclosed box on the right, with the time adjacent to the snapshots. PAR is shown in the licorice representation (silver) with its phosphate atoms (red) represented as spheres. Shown as spheres are $Mg^{2+}$ hexahydrates (blue), $Na^+$ (green) and $Cl^-$ (yellow) located within 6 Å of PAR.

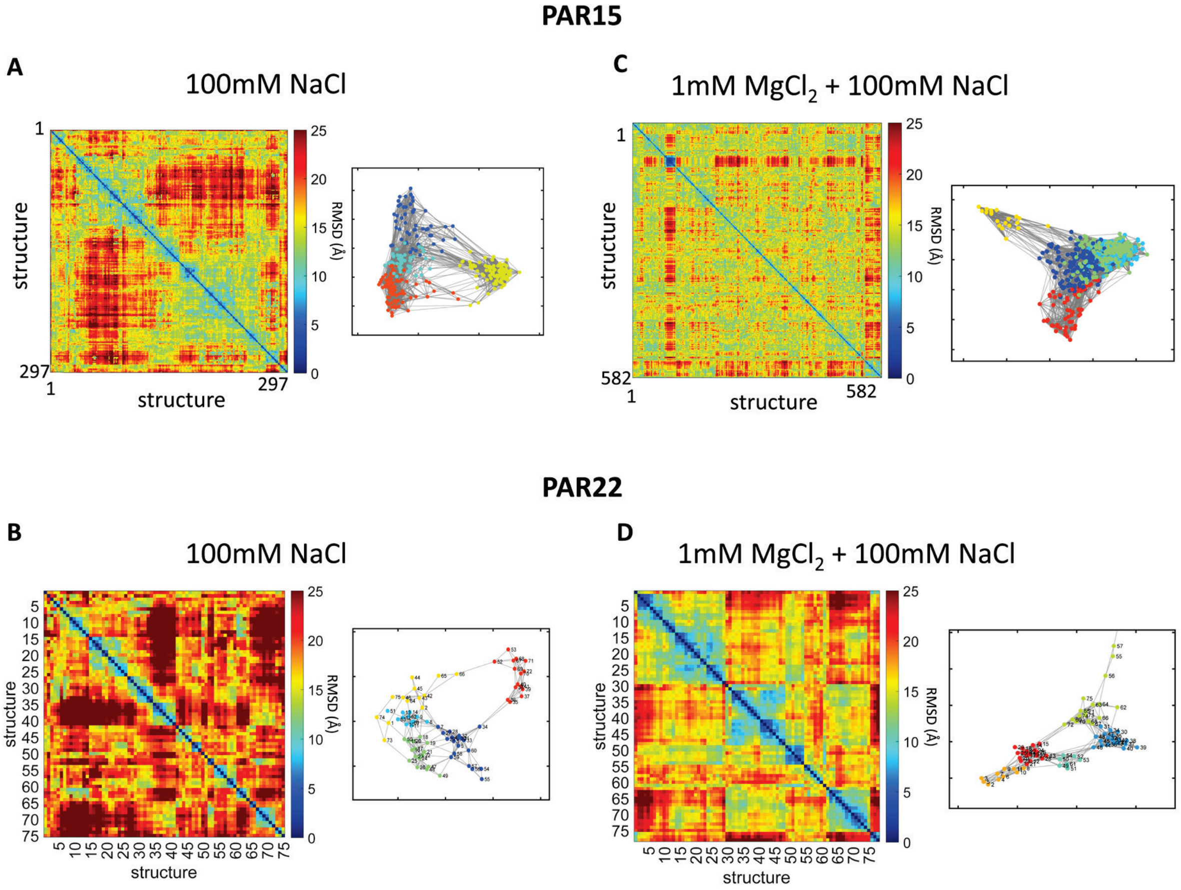


**Figure S10. Using graph theory to identify unique subclasses in PAR disordered structural ensembles.** For each experimental condition probed in this study, matrices of pairwise RMSD values are shown for $PAR_{15}$ and $PAR_{22}$. These matrices were transformed into binary adjacency matrices (not shown) by imposing a threshold RMSD value and spectrally clustered to map the structures into network graphs, where each node represents a structure and nodes are connected by an edge if their RMSD value falls below the threshold. Graphs are shown on the right of the RMSD matrices. The graphs were then clustered using the K-means approach to identify conformational subclasses. Different colors in each graph represent different K-means clusters, and the distance between nodes on the graph is representative of the distances in RMSD values between structures.

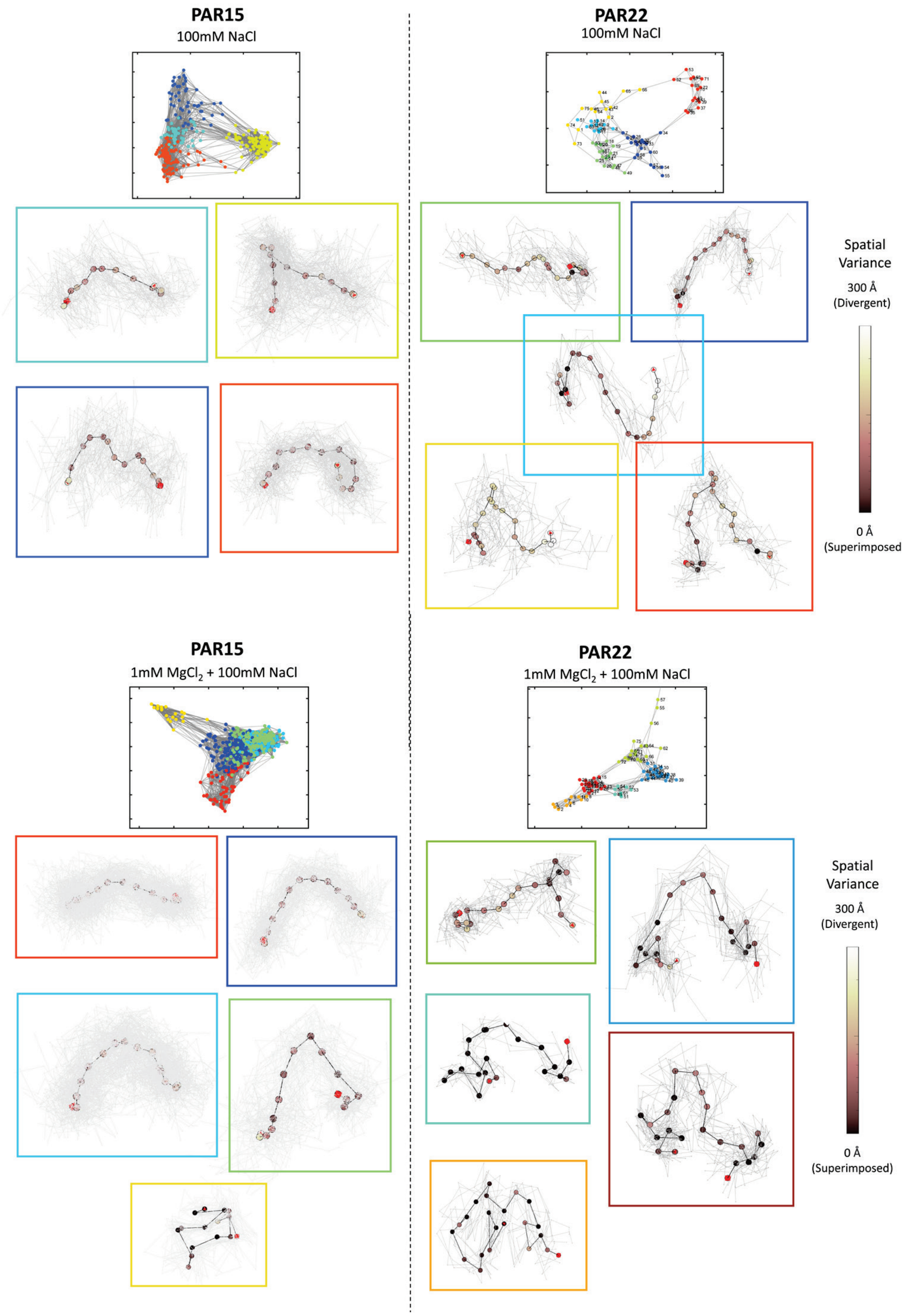


**Figure S11**. PAR structural classes plotted with all aligned structures overlaid atop the mean backbone conformation for visualization at the population level. Each point represents the mean position of every two phosphorus atoms along the backbone, colored by how much spatial variance is present across each subclass at that position. Cases are shown when only $Na^+$ is present at the top half of the figure and when both $Na^+$ and $Mg^{2+}$ are present at the bottom half. The different colored boxes around each subclass match the colors of their corresponding cluster within the network graph.
